# DNA Origami Voltage Sensors for Transmembrane Potentials with Single-Molecule Sensitivity

**DOI:** 10.1101/2021.08.18.456762

**Authors:** Sarah E. Ochmann, Himanshu Joshi, Ece Büber, Henri G. Franquelim, Pierre Stegemann, Barbara Saccà, Ulrich F. Keyser, Aleksei Aksimentiev, Philip Tinnefeld

## Abstract

Signal transmission in neurons goes along with changes in the transmembrane potential. To report them, different approaches including optical voltage-sensing dyes and genetically encoded voltage indicators have evolved. Here, we present a DNA nanotechnology-based system. Using DNA origami, we incorporate and optimize different properties such as membrane targeting and voltage sensing modularly. As a sensing unit, we use a hydrophobic red dye anchored to the membrane and an anionic green dye at the DNA connecting the DNA origami and the membrane dye anchor. Voltage-induced displacement of the anionic donor unit is read out by changes of Fluorescence Resonance Energy Transfer (FRET) of single sensors attached to liposomes. They show a FRET change of ∼5% for ΔΨ=100 mV and allow adapting the potential range of highest sensitivity. Further, the working mechanism is rationalized by molecular dynamics simulations. Our approach holds potential for the application as non-genetically encoded sensors at membranes.

## INTRODUCTION

The electrical transmembrane potential ΔΨ on the cellular level is a key parameter in the neurosciences. The introduction of fluorescence-based voltage sensors was a milestone towards a broader application and non-invasive visualization approaches in contrast to electrophysiological approaches being invasive, serial and time consuming.^1^ Many challenges with respect to signal, contrast and response time have been addressed with Genetically Encoded Voltage Indicators (GEVIs)^2–4^ that offer targetability to cell membranes. For improved contrast and imaging durations, hybrid approaches combining GEVIs with organic fluorophores have been introduced.^5,6^ These approaches, however, require transfected cell lines or transgenic animals.

In contrast, conventional voltage-sensing dyes face the challenge that all functionalities including targeting membranes, sensing and transducing a signal, have to be encoded in simple, chemically accessible structures. The development of a first generation of sensors yielded low-contrast Stark-effect voltage-sensing dyes and probes that disturbed cellular functions e.g. by capacitive loading of the membrane.^7^ Therefore, in recent approaches the complexity of sensors has been increased including bottom-up nanotechnological ideas to develop e.g. quantum-confined semiconductor nanoparticles or quantum dot-fullerene bioconjugates for voltage sensing.^8–10^ Recently, DNA was used as scaffolding material to combine electron-transfer based voltage-sensing dyes^7,11,12^ with targeting and intensity referencing for voltage sensing in organelles.^13^

Here, we use DNA origami to modularly address different challenges of voltage sensor design and demonstrate an alternative voltage-sensing strategy that allows sensing with bright dyes compatible with single-molecule imaging. DNA origami and similar self-assembly techniques offer the potential to meet the broad demands such as targeting lipid membranes, incorporating a sensing unit, providing a transduction mechanism optionally with internal referencing, being bio-compatible and minimally invasive.

In the DNA origami method, a long single-stranded DNA molecule (ssDNA, >7000 nucleotides long) is folded into a desired shape by hybridization with short oligonucleotides, producing billions of identical nanostructures.^14–16^ This bottom-up nanoassembly method offers the ability to place any chemical moiety by the integration of modified oligonucleotides on the nanostructure like on a molecular breadboard. Using the DNA origami technique, a variety of sensors has been realized,^17^ from nanopores^18–20^ to drug delivery systems^21,22^ to force sensors.^23,24^ By capturing DNA origami on nanocapillary tips, Hemmig and Fitzgerald *et al*. demonstrated the feasibility of using a DNA origami construct as a single-molecule voltage sensor.^25^ Two fluorophores capable of interacting *via* Fluorescence Resonance Energy Transfer (FRET) are placed on a DNA nanostructure such that, when subject to a voltage bias at the tip of a nanopipet, the FRET efficiency is modulated by the voltage magnitude.

Here, we demonstrate single-molecule transmembrane voltage read-out from the surface of a lipid membrane. By using a rectangular DNA origami for the arrangement of the different components needed, we create a sensor that optically reads out defined potentials via FRET with a change of ∼5% for ΔΨ=100 mV. FRET offers an advantageous ratiometric signal read-out and is therefore signal-intensity independent. We detect single FRET pairs by spacing out origami structures beyond the diffraction limit and hence, provide a pathway to image at the nanoscale beyond ensemble averages. We rationalize the functioning of the sensor through molecular dynamics (MD) simulations of the DNA-lipid membrane assembly. Further, we demonstrate the potential of DNA nanotechnology for voltage sensing by introducing small molecular changes in the sensing unit to shift the sensitivity of the sensor towards negative ΔΨ.

## RESULTS

The transmembrane voltage sensor is based on a rectangular DNA origami with dimensions of 70×100 nm^14,26,27^ functioning as a platform to program all the functionalities required into a small entity. In order to bind to liposomes, the nanostructure is equipped with ten cholesterol moieties and for binding to biotinylated PLL-e-PEG passivated surfaces, additional six biotin moieties are incorporated (Figure 1a, S1 and Table S1). Surface binding of the liposomes via the DNA origami facilitates imaging by total internal reflection microscopy (TIRF) while avoiding direct surface interactions of the liposomes.^28^ The voltage-sensing unit is placed centrally on the platform protruding from the structure. The hydrophobic and cationic dye ATTO647N connected to DNA by a C_12_-phosphate-C_6_ chain (C_12_+C_6_) is expected to anchor the sensor unit in the lipid membrane (Figure 1b and Figure S2 for molecular structures). Insertion into the lipid core of the membrane was observed for ATTO647N before.^29^ The DNA connection from ATTO647N to the DNA origami platform contains the anionic fluorophore ATTO532. We reason that any change of potential should have opposite effects on the average positions of the cationic ATTO647N dye and the anionic ATTO532 dye on the anionic DNA linker. The opposite forces on the two dyes should translate a ΔΨ into a change of FRET that can be read out optically and on the level of single molecules.

**Figure 1.**
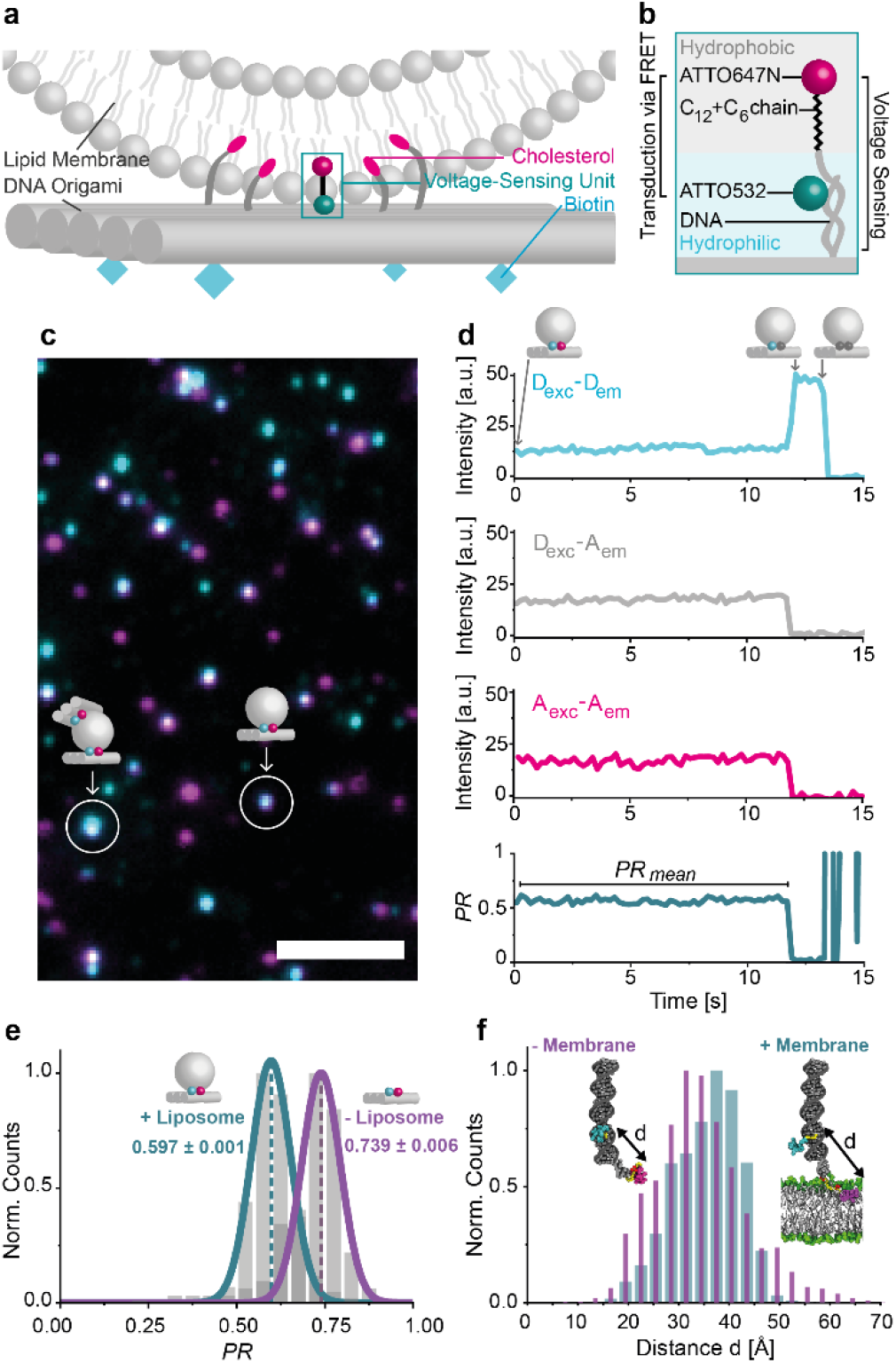
(a) DNA origami sensor of transmembrane potentials. A rectangular DNA origami plate is decorated with cholesterols to bind a liposome and with biotins to attach to the neutravidin-functionalized surface of a microscope’s cover slip. The voltage-sensing unit is positioned in the center of the DNA origami. (b) Voltage-sensing unit consisting of dsDNA protruding from the DNA origami plate and carrying an ATTO532 dye, and a complementary strand with an ATTO647N dye connected *via* a C_12_+C_6_ linker (see Figure S2 for chemical structure). The transduction of voltage signal to fluorescence is fulfilled by FRET from the donor ATTO532 to the acceptor ATTO647N. (c) Superimposed TIRF image of donor (blue) and acceptor (pink) fluorescence from DNA origami. White spots indicate DNA origami plates with both donor and acceptor dyes. The scale bar refers to 5 µm. (d) Single-molecule FRET transient. The fluorescence intensity over time is shown for the donor excitation-donor emission D_exc_-D_em_ channel (light blue), the donor excitation-acceptor emission D_exc_-A_em_ channel (grey) and the acceptor excitation-acceptor emission A_exc_-A_em_ channel (pink). From the D_exc_-D_em_ and the D_exc_-A_em_ channels, the *Proximity Ratio PR* and the *PR*_mean_ is determined (dark blue). (e) *PR* distributions for DNA origami constructs with (cyan) and without (purple) liposome attachment. The error refers to the standard error of the mean. *N*_molecule_ is ≥100 for each sample. (f) Histogram of inter-dye distances obtained from MD simulations of a dsDNA duplex decorated with the two dyes positioned at the lipid-water interface (cyan) and in aqueous solution (purple). Simulation times: - Membrane 1.35 µs, + Membrane 1.55 µs.

For imaging, we perform single-molecule FRET (smFRET) experiments^30^ of the optical potential sensor on a homebuilt TIRF microscope with green-red alternating laser excitation (ALEx, for details see SI).^31,32^ We acquire videos to follow the fluorescence over time and verify that single DNA origamis are observed. Figure 1c presents a superimposed TIRF image with donor dyes in cyan, acceptor dyes in pink and an overlay of the two in white. Some of the spots are brighter than others which is caused by multiple DNA origamis bound to a single liposome or origami multimers. To eliminate such aggregates in further analysis, we generate intensity-time transients from the videos for each spot with the software iSMS^33^ and inspect them visually. An exemplary transient is shown in Figure 1d with D_exc_-D_em_ (light blue), D_exc_-A_em_ (grey) and A_exc_-A_em_ (pink) with the subscript indicating the excitation and emission channels of donor (D) and acceptor (A), respectively. A correlated intensity increase in D_exc_-D_em_ upon an intensity decrease in D_exc_-A_em_ and A_exc_-A_em_ as well as a rapid photobleaching in D_exc_-D_em_ are a clear indication that indeed a single DNA origami is observed. From the intensities I_DD_ of the D_exc_-D_em_ and the intensity I_DA_ of the D_exc_-A_em_ channel, FRET is quantified as the *Proximity Ratio PR* with

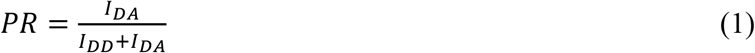

The *PR*_mean_ is calculated over the whole period of the energy transfer (bottom transient in Figure 1d) yielding one data point for each voltage sensor. All single-molecule transients are carefully reviewed and the ones showing a clear correlation between the three channels mentioned above are picked whereas transients showing multi-chromophore behavior are rejected.

We first test whether an interaction between the voltage sensor and the lipid membrane as intended is detected. To this end, we study the DNA origamis with and without 100 nm DOPC liposomes by mixing the origami structures with an excess of liposomes and immobilizing the complexes on a surface. After performing smFRET measurements, we obtain *PR* distributions as shown in Figure 1e. The liposome-free sample yields a mean *PR* of 0.739±0.006 (standard error of the mean, SEM) obtained from Gaussian fitting of the distribution which decreases to 0.597±0.001 for the liposome-containing sample (Figure 1e). The fact that we obtain narrow homogenous populations that are clearly shifted with respect to each other indicates quantitative binding of DNA origami voltage sensors to the liposomes. In addition, the decrease of FRET supports the idea that the hydrophobic ATTO647N is diving into the membrane core so that the average distance of donor and acceptor is substantially increased upon membrane binding.

We further rationalize the idea of the FRET-acceptor anchoring in the membrane by MD simulations of the voltage-sensing unit with and without a lipid membrane present (Figure 1f and S3). The Movie S1 provided in the SI reveals a coiling of the ATTO647N with the alkyl chain resulting in a close proximity of the dyes. This secondary structure is broken in the presence of a lipid membrane (Movie S2) as ATTO647N and the alkyl chain insert to and remain in the hydrophobic core of the membrane. Figure 1f shows the distributions for the inter-dye distances for both samples determined from the MD simulations. The observed shift towards larger distances for the sensor in presence of a membrane is in good agreement with the experimental results (Figure 1e) and suggests that the lower *PR* upon liposome addition is a result of the spatial separation of the two dyes. Another interesting observation from the simulations is that the ATTO647N dye remains embedded in the nearest to the membrane, interacting with the phosphate moieties of the lipid head groups (likely because of its positive charge) while the main body of the dye resides inside the hydrophobic core of the membrane.

In order to test the performance of our voltage-sensing DNA origami, we use ion exchange by the ionophore valinomycin^34^ to create a well-defined drop of ΔΨ across the liposome membrane. In a typical experiment, the origami-liposome complexes are imaged, the buffer surrounding is exchanged to introduce a potassium gradient across the lipid membrane and valinomycin is added, before the sample is imaged again (Figure 2a). Valinomycin specifically complexes potassium but not sodium ions and shuttles them across the lipid membrane until an equilibrium is reached and a polarized membrane results following the Nernst equation

**Figure 2.**
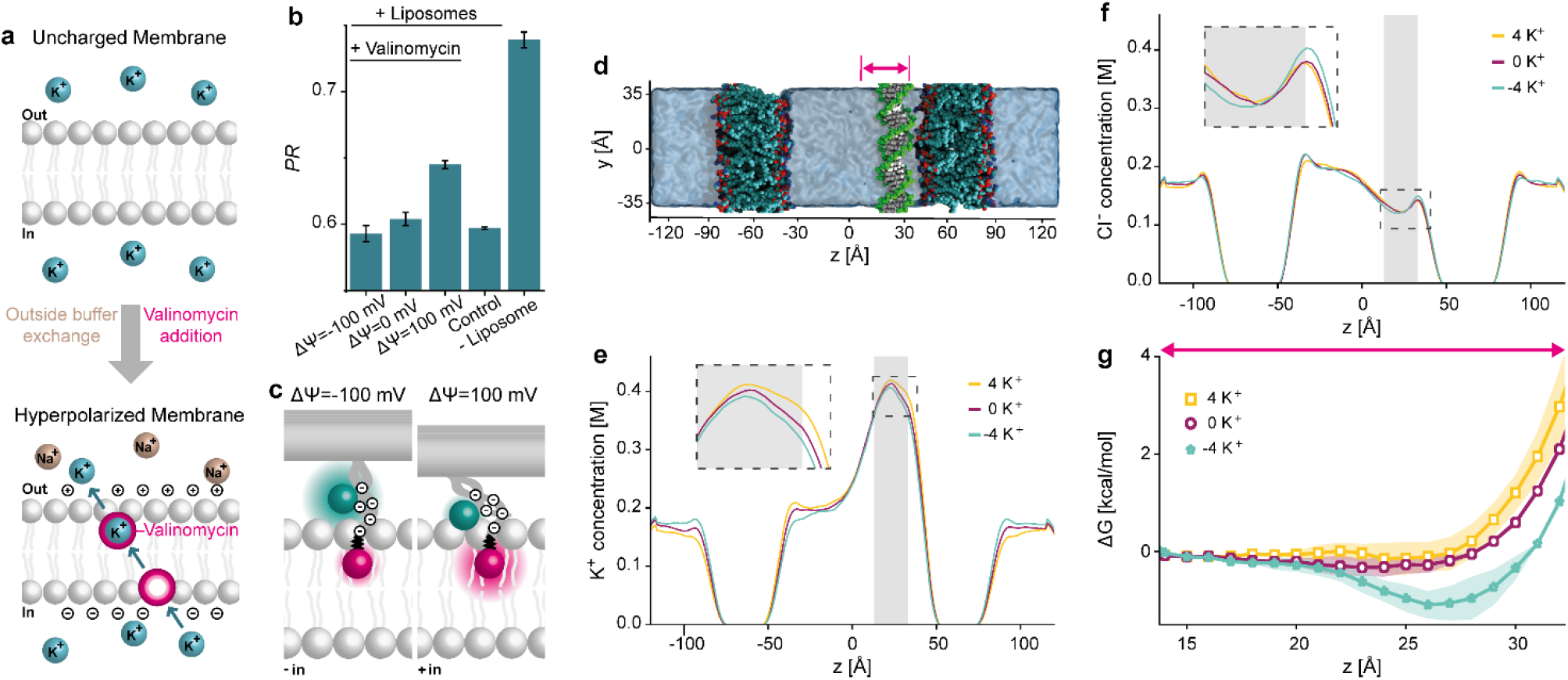
(a) Creation of electrical transmembrane potentials ΔΨ. By exchanging the outside buffer, a potassium ion gradient across the lipid membrane is built up. Equilibration of the potassium gradient by the ionophore valinomycin converts the chemical potential into an electrical transmembrane potential ΔΨ. (b) Mean *PR* and standard errors of the mean derived from Gauss fits to the distributions (Figure S5 and S6) of the DNA origami-liposome complexes with different ΔΨ in comparison to control samples presented in Figure 1e. *N*_molecule_ is ≥100 for each sample. (c) Proposed working principle of voltage-sensing DNA origami. The ATTO647N remains as an anchor in the membrane’s hydrophobic core whereas the surrounding DNA with its anionic nature is attracted towards the membrane by K^+^ excess inside of the liposome resulting in a shorter inter-dye distance and an increased FRET. (d) Representative configuration of a simulated double membrane system, where two membrane patches separate two compartments filled with 150 mM KCl solution. A single dsDNA molecule is placed near one membrane to characterize effective interactions between the DNA and the membrane. A gradient of K^+^ concentration is established by transferring four K^+^ ions from one compartment to the other, corresponding to a drop of ΔΨ=±1.3 V. The local concentration of K^+^ (e) and Cl^-^ (f) ions along the lipid bilayer is shown for the three ion gradient conditions. The z axis is defined in panel d. The profiles were averaged over 21 replica windows of the respective REUS MD simulations, each replica simulation being 120 ns long. The shaded region shows the location of the center of DNA in various windows. (g) Free energy ΔG of the 21 base pair dsDNA as a function of its *z*-coordinate for the three ion gradient conditions. The arrow implies the region shown in (d).

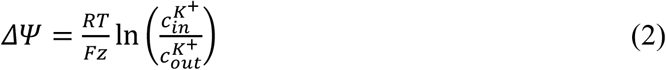

where R is the gas constant, T is the temperature, F is the Faraday constant, z is the charge number and 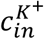 and 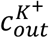 are the potassium concentrations inside and outside of the liposome, respectively. By adjusting the initial K^+^ concentration gradient, we produce well-defined ΔΨ (Table S2). Figure S4 confirms the functionality of our assay in bulk experiments using a commercially available voltage-sensing dye.

First, we are interested in three scenarios: a hyperpolarized, a neutral and a depolarized membrane with respect to the inner leaflet. We choose the hyperpolarization to be ΔΨ=-100 mV and the depolarization to be ΔΨ=100 mV, for which the buffer outside is exchanged with respect to the desired ΔΨ and valinomycin is added before imaging. For all the samples, single Gaussian distributions are obtained and mean *PR* values of 0.593±0.006 for ΔΨ=-100 mV, of 0.604±0.005 for ΔΨ=0 mV and of 0.645±0.003 for ΔΨ=100 mV are determined (Figure 2b and S5). When compared to the liposome-free sample, all the *PR* values are lower, which, in combination with the mono-Gaussian nature of the distributions, strongly suggests that the liposomes stayed intact throughout the experimental procedure. As the ΔΨ=0 mV sample shows an almost identical *PR* histogram as the control sample before the valinomycin addition, we are confident that all observed changes in the single-molecule fluorescence result from the ΔΨ created and are not an interference with the ionophore (Figures 2b and S6). In contrast, there is a notable increase in the *PR* for the depolarized membrane compared to the hyperpolarized and neutral membrane which implies that the DNA origami-based sensor is able to report transmembrane potentials on the single-molecule level.

The direction of FRET change implies that a more positive charge on the inside would attract the anionic donor dye-DNA hybrid towards the membrane so that FRET increases (see Figure 2c). An alternative mechanism, where the change of ion concentration outside the membranes modules the electrostatic force acting on the dye embedded in the lipid membrane is ruled out through a set of MD simulations that examined the distribution of the electrostatic potential in a double membrane system (Figure S7 and S8). It has been previously established that the electric potential of the membrane’s interior is approximately 500 mV higher than the electric potential of the surrounding electrolyte.^35,36^ A slight imbalance of ion concentration, i.e., a transfer of just one ion between the compartments of our simulated double membrane system as shown in Figure S7 produces the expected voltage difference between the electrolyte compartments. However, the gradient of the electrostatic potential across the leaflets of the lipid bilayers remains largely unaffected by the ion concentration gradient as most of the additional potential drop occurs at the interface of the lipid head groups and the electrolyte which is why we rule out a movement of the membrane-anchored ATTO647N.

To directly probe the effect of an ion concentration gradient on the interaction between DNA and a lipid membrane, we simulate another double membrane system (Figure 2d) where one DNA molecule is placed near the surface of one of the membranes parallel to the membrane surface. In addition to the system containing two charge-neutral electrolyte compartments (0 K^+^), two variants of the system are created by moving four K^+^ either to (4 K^+^) or from (−4 K^+^) the compartment containing the DNA, which corresponds to ΔΨ=0, ΔΨ=+1.3 and ΔΨ=-1.3 V, respectively (Figure S8). Such higher than experimental bias conditions are chosen to increase the effective force on dsDNA, facilitating convergence of the free-energy calculations (describe below). Replica exchange umbrella sampling (REUS) simulations^37^ are performed for each system using 21 sampling windows (in 1 Å increments) for the distance between the centers of mass of the dsDNA and the nearby membrane along the *z*-axis. The resulting ion gradient produces the expected ΔΨ across the compartments (Figure S8). Further, the local concentration of K^+^ (Figure 2e) and Cl^-^ (Figure 2f) ions show a non-trivial behavior. In the profiles for all three samples, it is clearly visible that the K^+^ concentration is increased close to the DNA whereas the Cl^-^ concentration is decreased which is due to the electrostatic attraction or repulsion to the anionic DNA backbone, respectively. In the case of an excess of K^+^ ions inside (4 K^+^), the K^+^ concentration is also higher close to the inner leaflets. Interestingly, the concentration at the respective outer leaflets is reduced indicating a capacitive effect. The opposite behavior is observed for the lack of K^+^ ions inside (−4 K^+^). A complementary effect is observed for the Cl^-^ concentration (Figure 2f).

Further analysis of the REUS simulations yields the free energy of the dsDNA as a function of its proximity to the lipid membrane (Figure 2g). In the absence of a K^+^ gradient, the free energy has a shallow minimum near the membrane surface, in agreement with our previous calculations.^38^ Moving the positive charge across the membrane from the compartment housing (−4 K^+^ trace), produces a free-energy minimum near the membrane surface promoting DNA attraction to the membrane surface. Moving the positive charge into the DNA compartment (4 K^+^ trace) slightly increases repulsive interaction between DNA and the lipid membrane. These simulation results in qualitative agreement with our observation of a FRET increase for depolarized membranes and support the mechanism shown in Figure 2c.

Next, we study the sensitivity of our voltage sensor in more detail and vary the potentials from ΔΨ=-125 mV to ΔΨ=125 mV in steps of 25 mV. For each sample the mean *PR* before creating ΔΨ is approximately the same (Figure S9). We therefore merge all reference data and define it as the mean of the control sample *PR*_*before*_. This value is subtracted from the *PR* after ΔΨ is built up (Figure S5) as

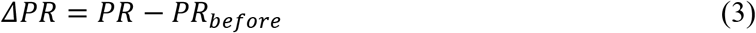

yielding the change *ΔPR*. The respective SEM is derived after Gaussian error propagation (see SI) and the data is presented in Figure 3. In accordance with the results discussed above, the *PR* value is only slightly increasing up to ΔΨ=50 mV and increases strongly in the range of 50 to 100 mV. The voltage sensor is thus able to transduce small changes in ΔΨ to single-molecule fluorescence signals. The non-linear response might indicate that the sensing unit above the membrane is not progressively shifting in the changing ΔΨ but that more specific conformational changes or displacements of the dyes are occurring.

**Figure 3.**
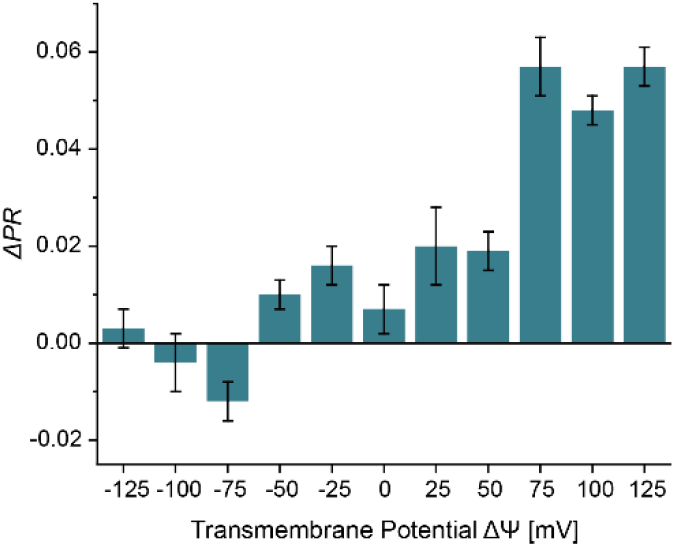
Changes *ΔPR* of the voltage sensor exposed to liposomes with different electrical transmembrane potentials ΔΨ. *ΔPR* is calculated by subtracting the mean *PR* before the potentials ΔΨ is created from the respective *PR* of the sample as indicated. The error bars represent the standard error of the mean after Gaussian error propagation. *N*_molecule_=100 for each sample.

As our proposed mechanism is strongly relying on the relative positioning of the donor dye with respect to the acceptor dye, we check the sensitivity of the system for small changes of the linker. We therefor change the voltage-sensing unit minimally by shortening the carbon chain from a C_12_+C_6_ to a C_12_ chain eliminating also the additional phosphate group (Figure 4a, for details see Figure S2). Interestingly, in the absence of the liposomes, the *PR* is only minimally higher for the shorter linker (*PR*=0.754 instead of 0.739). Upon binding to the liposome, however, the *PR* does only slightly decrease to *PR*=0.732 for the shorter linker indicating that stretching of the hydrophobic linker is mainly responsible for the FRET reduction in case of the C_12_+C_6_ linker (Figure S10 and S11). Varying ΔΨ of the liposomes exposed to the DNA origami voltage sensor with the shortened linker has an interesting effect on the measured *PR* values. Most of the signal change now occurs in the more physiologically relevant range between -100 mV and 0 mV whereas only a small *PR* increase is detected for positive ΔΨ (Figure 4b, S11 and S12). The direction of change is compatible with the idea that the FRET reduction is not a linear displacement in the ΔΨ but related to a more specific conformational change. As the DNA and the negatively charged dye are pulled towards the membrane by the shorter linker, it requires a more negative potential to displace them from the membrane so that the FRET reduction occurs.

**Figure 4.**
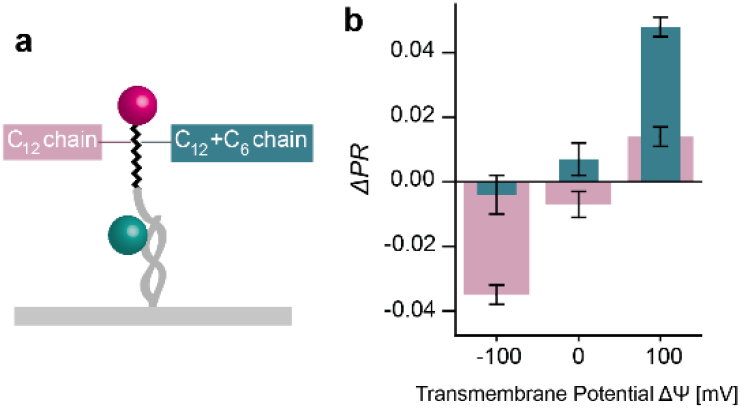
Changes *ΔPR* of the sensor with a C_12_ chain in the voltage-sensing unit as illustrated in (a) exposed to liposomes with ΔΨ=-100 mV, ΔΨ=0 mV and ΔΨ=100 mV (b, pink) in comparison to the voltage sensor with a C_12_+C_6_ chain (blue). *ΔPR* is estimated by subtraction of the mean *PR* before the potentials are created from the respective *PR* of the sample as indicated. The error bars represent the standard error of the mean after Gaussian error propagation. *N*_molecule_ is ≥91 for each sample.

## CONCLUSION

Transmembrane potentials are key parameters to understand cellular functions and interactions and there is a great need for the development of smart sensing systems. We here present a DNA origami voltage sensor offering a robust platform to include many functionalities such as surface immobilization and liposome binding. It could be extended for further smart functionalities including specific cell or organelle recognition or for immune system camouflage.^39–41^

We also introduce a new sensing unit that is based on FRET between a hydrophobic dye that prefers location in the hydrophobic membrane core and a hydrophilic and anionic dye-DNA moiety that reacts with a *PR* change of ∼5% for ΔΨ=100 mV. The DNA origami voltage sensors are studied by single-molecule spectroscopy on liposomes and the results are rationalized by MD simulations. While the fundamental working principle is implied by the experimental results, the MD simulations provide evidence that more specific interactions between the membrane and the sensing unit determine the sensitive voltage range that could be tuned by adaptation of the linker between donor and acceptor. Overall, our data show profound potential for this novel approach for ΔΨ sensors that could similarly be adapted for other sorts of sensors.

## ASSOCIATED CONTENT

### Supporting Information

Supporting Information: Material and Methods, Illustration of DNA origami design in CaDNAno, DNA oligonucleotides used as staple strands for the DNA origami voltage sensor, Detailed sketch of the different voltage-sensor designs, Equilibrium MD simulation of dye conjugated dsDNA in aqueous and membrane anchored environment, Concentration of KCl and NaCl in the buffer inside and outside of the LUVs, Valinomycin bulk test, *PR* distributions for all samples of the C_12_+C_6_ sensor with transmembrane potentials ΔΨ, *PR* distributions of various C_12_+C_6_ samples compared, Voltage bias created by shuffling of a single ion in double membrane systems, Average electrostatic profile of the 0 K^+^, 4 K^+^ and -4 K^+^ systems along the bilayer normal during the REUS MD simulations, *PR* distributions for all control samples of the C_12_+C_6_ sensor, *PR* distributions for all control samples of the C_12_ sensor, *PR* distributions for samples of the C_12_ sensor with transmembrane potentials ΔΨ and without liposomes, *PR* distributions of various C_12_ samples compared, Captions to Supplementary Movies (docx)

Movie showing all-atom MD simulation of dyes on dsDNA (MP4)

Movie showing all-atom MD simulation of dyes on dsDNA with lipid membrane (MP4)

Movie showing mrDNA simulation of the DNA origami (MP4)

## AUTHOR INFORMATION

### Notes

The authors declare no competing financial interest.

## ACKNOWLEDGMENT

This work has been supported by the Deutsche Forschungsgemeinschaft DFG (grants INST 86/1904-1 FUGG, TI 329/10-1 and Project - ID 201269156 - SFB1032), and the National Science Foundation (USA) (DMR-1827346). We acknowledge the supercomputer time provided to A.A. and H.J. through the XSEDE allocation grant (MCA05S028) and the Leadership Resource Allocation (MCB20012) on Frontera at Texas Advanced Computing Center. A.A. and H.J. would like to thank Christopher Maffeo for his help in setting up the initial mrDNA simulation.

## ABBREVIATIONS

GEVI: Genetically Encoded Voltage Indicator;
FRET: Fluorescence Resonance Energy Transfer;
ssDNA: single-stranded DNA;
MD: Molecular Dynamic;
TIRF: Total Internal Reflection Fluorescence;
dsDNA: double-stranded DNA;
smFRET: single-molecule FRET;
ALEx: Alternating Laser Excitation;
*PR*: *Proximity Ratio*;
SEM: Standard Error of the Mean;
REUS: Replica Exchange Umbrella Sampling.

## SYNOPSIS

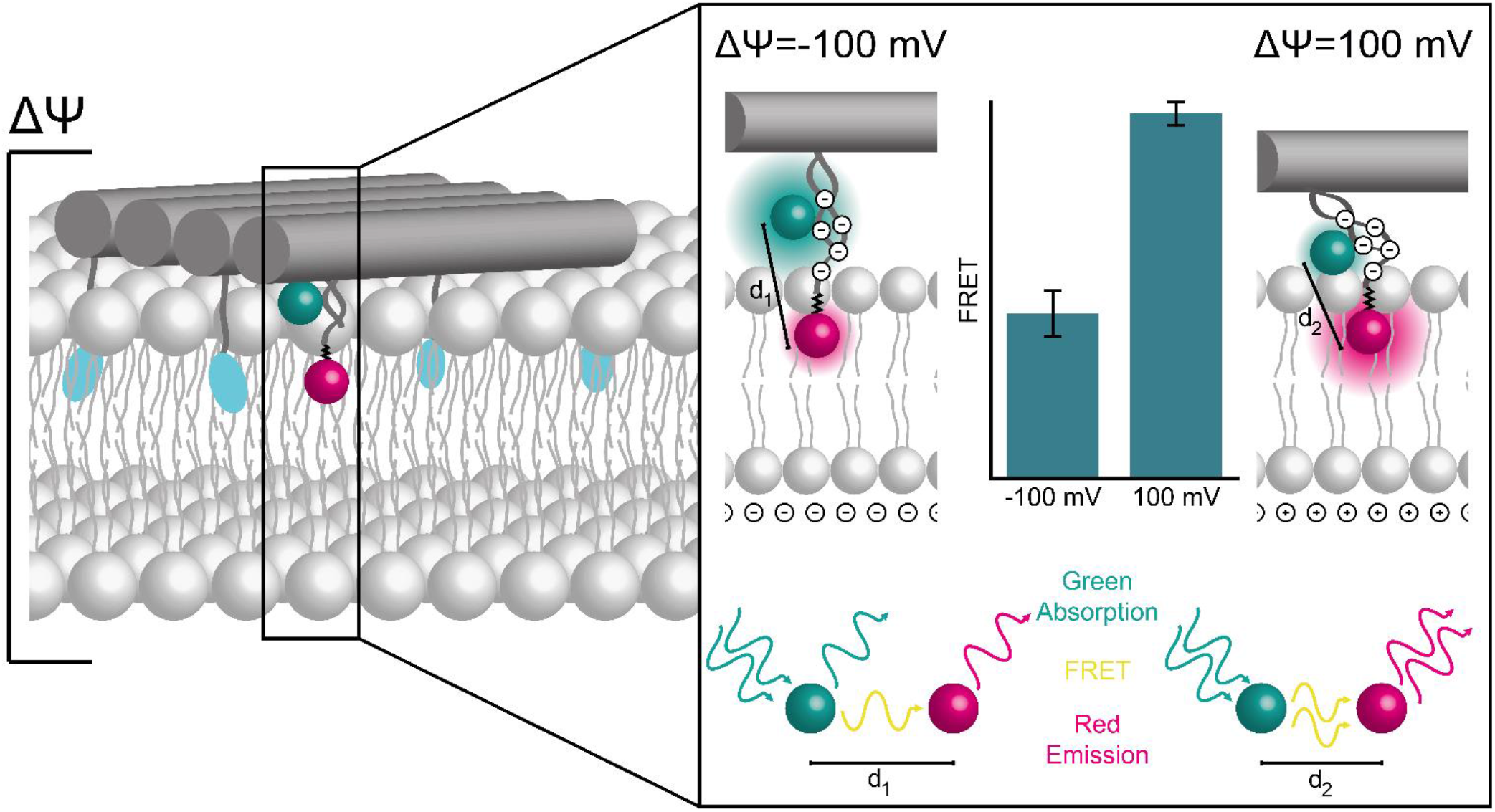

