## Supporting Information for "DNA Origami Voltage Sensors for Transmembrane Potentials with Single-Molecule Sensitivity"

### Material and Methods

#### *Experimental part*

##### Chemicals

If not declared differently, all chemicals were purchased from Merck KGaA.

##### DNA origami folding

A rectangular DNA origami[1] was used which is based on a 7249 nt long scaffold derived from the M13mp18 bacteriophage. It was designed in the software CaDNAno[2] and carried several modifications as indicated in Figure S1. All staple strand sequences are given in Table S1 together with the name of the company from which it was purchased.

The origami structures were folded in buffer containing 10 mM Tris, 1 mM EDTA and 12.5 mM $MgCl_2$ by mixing 10 nM of the scaffold with 100 nM of unmodified and 300 nM of modified oligonucleotides. The solution was heated to 70°C for 3 min and then cooled down in 1°C-steps remaining 1 min at each temperature in a thermocycler (primus 25, peqlab). During the folding process, the biotin modified oligonucleotides, the binding site containing an ATTO532 dye and three cholesterol strands on a ssDNA leash were incorporated. By PEG precipitation, the folded structures were purified from excess staple strands where the sample was mixed in a 1:1 ratio with a buffer containing 12% PEG-8000 (w/v), 10 mM Tris, 1 mM EDTA, 500 mM NaCl and 12 mM $MgCl_2$ at pH 7.5 and centrifuged for 30 min at 16 krcf and 4°C. The supernatant was discarded and the pellet dissolved in the buffer used for folding. This step was repeated 4 times. In order to label the DNA origami with seven more cholesterols and the voltage-sensing unit, it was incubated over night at room temperature with 5x excess of the oligonucleotides named above per binding site, before another PEG precipitation for purification was examined. The samples were stored at 4°C until usage.

##### Liposome preparation

To produce Large Unilamellar Vesicles (LUVs), lipid films were created. Therefor 1,2-dioleoyl-sn-glycero-3-phosphocholine (DOPC, Avanti Polar Lipids, INC.) was dissolved at a concentration of 25 mg/mL in chloroform and 1 mmol was added to a glass vial, dried under a nitrogen stream and for another 4 h under vacuum in a desiccator. These lipid films were stored at -20°C until further usage. To create LUVs, the lipid films were dissolved in LUV buffer containing 5 mM Tris, 1 mM EDTA, 0.5 mM Trolox and either 150 mM KCl or 149 mM NaCl and 1 mM KCl at pH 7 (potassium or sodium LUV buffer, respectively) resulting in a lipid concentration of 2.5 mM. After seven freeze-and-thaw cycles using liquid nitrogen and a 80°C water bath, the solution was extruded with a LiposoFast Basic extruder (Avestin, INC.) using Nucleopore PC membranes with a pore size of 100 nm (Whatman, Cytiva Ltd.).

##### Preparation of microscope slides

Nunc Lab-Tek II Chambered Slides (Thermo Fisher Scientific Inc.) were cleaned with 1 M KOH for 4 h at room temperature. After washing with 1xPBS buffer, the slides were passivated over night at 4°C with 0.5 mg/mL PLL(20)-g[3.5]- PEG(2)/PEG(3.4)- biotin(50%) (PLL-PEG-biotin, SuSoS AG) in 1xPBS. After washing with 1xPBS, 0.25 mg/mL NeutrAvidin (Thermo Fisher Scientific Inc.) in 1xPBS was added for 20 min and washed off with 1xPBS and the slides were ready to use.

#### TIRF microscope

A homebuilt Total Internal Reflection (TIRF) microscope based on an Olympus IX71 inverted microscope was used for the single-molecule Fluorescence Resonance Energy Transfer (smFRET) measurements. The beams of a green laser (Sapphire 532 nm, 100 mW, Coherent) and a red laser (iBeam Smart 640 nm, 150 mW, Toptica Photonics) were altered using an acousto-optical tunable filter (AOTF, PCAOM-VIS, Crystal Technology) at a frequency of 10 Hz. The power of the green laser was set to 30 mW and of the red laser to 80 mW. The light was focused on the sample with an oil-immersion objective (APO N 60XO/ 1.49 NA TIRF, Olympus). The emission light was separated from the excitation light by a dual line beamsplitter and further separated into 2 emission channels with an Optosplit III (Cairn Research) equipped with a dichroic beam splitter (640 DCXR, Chroma Technology). The green emission was spectrally filtered with a bandpass filter (BrightLine HC 582/75, Semrock) and the red emission with a longpass filter (647 nm RazorEdge, Semrock), before being focused on a back-illuminated sCMOS camera (KURO 1200B sCMOS, Princeton Instruments) in a dual-view configuration. The LightField software (Princeton Instruments) was used to acquire videos with a length of each min. 300 frames.

#### Sample preparation and imaging

To obtain DNA origami-liposome complexes, the DNA origami were incubated with the liposomes with a 100x excess of the liposomes over the origami for 2 h at room temperature in the respective LUV buffer (potassium or sodium). Then the structures were immobilized in Lab-Tek chambers at a concentration of 30 pM via biotin-neutravidin interaction and imaged on the homebuilt TIRF microscope described above. The LUV buffer matching the ion composition inside of the liposome was used. These measurements represent the control samples, before in each of these samples an electrical transmembrane potential ΔΨ was built up.

To then create the electrical transmembrane potential ΔΨ, first the buffer surrounding the origami-liposome complexes was exchanged with respect to the desired potential. To determine the ionic gradient required, the Nernst equation

$$\Delta\Psi = \frac{RT}{Fz} ln\left(\frac{c_{in}^{K^+}}{c_{out}^{K^+}}\right) \quad (1)$$

with R as the universal gas constant, T as the temperature, F as the Faraday constant, z as the charge number and $c_{in}^{K^+}$ or $c_{out}^{K^+}$ as the $K^+$ concentration inside or outside of the liposome, respectively, was used. Table S2 shows the concentrations of KCl and NaCl used to create the different electrical potentials tested. Secondly, 7.5 nmol of valinomycin was added which locates into the hydrophobic core of the lipid bilayer, complexes potassium ions and shuttles them through the lipid membrane while the Chloride counter ions remain. Thereby an electrical transmembrane potential ΔΨ is built up.[3] After an incubation of 10 min with the valinomycin, the samples were imaged again on the homebuilt TIRF microscope. For the liposome-free sample, the DNA origami were immobilized without prior LUV incubation and imaged in buffer containing 10 mM Tris, 1 mM EDTA, 0.5 mM Trolox, 149 mM NaCl and 1 mM KCl at pH 7.

#### Data analysis

For the data analysis, the software iSMS[4] running on Matlab was used. The split channels of green and red emission were superimposed and from the videos an intensity-time transient was derived for each single spot. These transients were then carefully revised to separate single DNA origami structures showing FRET from multimers or origami not containing both fluorophores. In the three channels of donor excitation-donor emission $D_{exc}$-$D_{em}$, donor excitation-acceptor emission $D_{exc}$-$A_{em}$ and acceptor excitation-acceptor emission $A_{exc}$-$A_{em}$ it was checked for a correlation typical for single-molecule FRET pairs. If there was a clear correlation between the different channels – e.g. an intensity increase in the $D_{exc}$-$D_{em}$ channel while a decrease in the $D_{exc}$-$A_{em}$ channel is observed upon a simultaneous drop in the $A_{exc}$-$A_{em}$ channel – the transient was picked and the period selected over which the mean *Proximity Ratio PR* was calculated as

$$PR = \frac{I_{DA}}{I_{DD}+I_{DA}} \tag{2}$$

with $I_{DA}$ as the intensity from the $D_{exc}$-$A_{em}$ channel and $I_{DD}$ as the intensity from the $D_{exc}$-$D_{em}$ channel. The data derived this way was further plotted against its frequency and a Gauss fit was used to determine the mean *PR* and its standard error for each sample.

The change *ΔPR* shown in Figure 3 and 4 was calculated as

$$\Delta PR = PR - PR_{before} \tag{3}$$

with *PR* as the value determined for the respective sample with the transmembrane potential $\Delta\Psi = x\,mV$ and $PR_{before}$ as the value derived before the addition of valinomycin. The respective standard error $\sigma_{\Delta PR}$ resulted from a Gaussian error propagation as

$$\sigma_{\Delta PR} = \sqrt{\left(\left|\frac{\partial \Delta PR}{\partial PR}\right| \cdot \sigma_{PR}\right)^2 + \left(\left|\frac{\partial \Delta PR}{\partial PR_{before}}\right| \cdot \sigma_{PR_{before}}\right)^2} \tag{4}$$

#### Valinomycin bulk test

To proof that valinomycin creates electrical transmembrane potential ΔΨ in liposomes that have a potassium gradient between the in- and outside, a bulk assay was performed using the voltage-sensitive fluorophore 3,3'-Dipropylthiadicarbocyanine Iodide ($DiSC_3(5)$, Thermo Fisher Inc.), liposomes and the spectrofluorometer FS5 (Edinburgh Instruments). The $DiSC_3(5)$ dye is cationic and accumulates on hyperpolarized lipid membranes where its fluorescence is reduced due to contact quenching. Hence, the fluorescence intensity depends on the electrical potential of the lipid membrane.

High precision cell cuvettes (Ultra-Micro Cell 105.252-QS, Hellma analytics) were passivated with a 1 mg/mL BSA solution (Sigma Aldrich) in 1xPBS buffer to decrease unspecific binding. 100 µL of LUVs with a lipid concentration of 200 µM were added. The buffer inside the liposome for each sample tested contained 5 mM Tris, 1 mM EDTA, 0.5 mM Trolox and 150 mM KCl at pH7. The buffer in which the LUVs were diluted to the respective concentration was either the same or instead of the 150 mM KCl contained 149 mM NaCl and 1 mM KCl. The respective buffer combination for each sample is depicted in Figure S4.

The $DiSC_3(5)$ dye was added to the liposome containing cuvette resulting at a final concentration of 1 μM and an addition of 1% (v/v) DMSO to the solution. After an incubation of 10 min, the sample was placed in the spectrofluorometer and the acquisition started ($\lambda_{ex}$=666 nm, $\text{bandwidth}_{ex}$=1 nm, $\lambda_{em}$=691 nm, $\text{bandwidth}_{em}$=5 nm, 1 point/s). Once the fluorescence intensity was stable, valinomycin was added resulting in a final concentration of 500 nM and another 0.25% (v/v) DMSO, so the final DMSO concentration in the cuvette was 1.25% (v/v). Then the effect of valinomycin was followed by tracking of the fluorescence intensity. For the samples testing the influence of the buffer on the fluorescence intensity, instead of the valinomycin solution, only the respective buffer outside of the liposome was added together with the same overall DMSO concentration of 1.25% (v/v).

To first neglect an interference between the $DiSC_3(5)$ and valinomycin, the ionophore was added to the free dye (Figure S4a). After the addition, the fluorescence baseline is lower than before, but no further effect is observed. The intensity drop can be affiliated to unspecific binding of the dye to the pipette tip; the overall dye concentration is slightly reduced and as a consequence, the fluorescence signal, too. This underlines the importance to not mix the following samples by multiple pipetting to reduce the effect.

Next, the effect of adding a solution to the liposome-dye mixture was investigated. Therefor the respective buffer with DMSO was added to liposomes with inside and outside potassium buffer (Figure S4b) and with inside potassium and outside sodium buffer (Figure S4c) liposomes. After the addition, for both samples an equilibration towards a higher intensity is observed which is related to a homogeneous distribution after Brownian motion in the field of view. As for the free dye, the baseline is slightly reduced though due to unspecific dye sticking to the pipette tip. Anyhow, no specific intensity decrease is observed. Next, the effect of valinomycin to the liposome-dye mixture was tested when there is no ion gradient to the outside (Figure S4d) and as previously, no change is observed which means that no polarization at the lipid membrane is induced.

Lastly, as in the single molecule experiments, valinomycin was added to a sample with potassium carrying liposomes in sodium buffer with $DiSC_3(5)$ (Figure S4e). Unlike in the samples before, after the valinomycin addition the intensity baseline is not only lower, but also an equilibration towards a lower intensity is observed. This different behavior clearly proofs that a transmembrane potential $\Delta\Psi$ is built up and the cationic voltage-sensing dye is attracted to the hyperpolarized membrane where its accumulation leads to contact quenching and hence, a reduced overall intensity.

#### *Simulation part*

##### General simulation protocols

All MD simulations were performed using program NAMD2[5], a 2 fs integration time step, 2-2-6 multiple time stepping, periodic boundary conditions and particle mesh Ewald (PME) method over a 1 Å resolution grid to calculate the long range electrostatic interaction.[6] The Nose-Hoover Langevin piston[7] and Langevin thermostat were used to maintain the constant pressure and temperature in the system. An 8-10-12 Å cutoff scheme was used to calculate van der Waals and short-range electrostatic forces. SETTLE algorithm[8] was applied to keep water molecules rigid whereas RATTLE algorithm[9] constrained all other covalent bonds involving hydrogen atoms. CHARMM36 force field parameters described the bonded and non-bonded interactions among the atoms of DNA[10], lipid[11], water and ions.[12] Magnesium ions were modeled as magnesium hexahydrates ($Mg[H_2O]_6^{2+}$).[13] Corrections to non-bonded interactions potentials were applied to improve description of ion-DNA9, ion-ion, and DNA-lipid interactions.[14] CHARMM General Force Field (CGenFF)[15] were used to describe the interaction parameters for the dye molecules. The coordinates of the system were saved at an interval of 20 ps. The visualization, analysis and post-processing of the simulation trajectories were performed using VMD[16] and CPPTRAJ.[17]

#### Initial models of lipid bilayer membrane and dye conjugated DNA

Starting with the caDNAno design of DNA origami plate along with the modified strands for anchoring and voltage sensing (Figure S1 and Table S1), we performed coarse-grained MD simulation using mrDNA resulting in the movie S3.[18] The final configuration of the origami plate at the end of the coarse-grained simulation was converted to an all-atom model. In order to obtain the nanoscale structure and dynamics of the dye molecules conjugated to the DNA fragment, we selected the 22 base-pair long DNA strand containing the dye molecules in experimental design. The topology and parameters file for the ATTO647N and ATTO532 dye molecules covalently conjugated using $C_{12}$+$C_6$ linker molecules to DNA were obtained using the CHARMM General Force Field (CGenFF) webserver.[19] We used a custom psfgen script in VMD to covalently connect the dye molecules to DNA according to the chemical sketch shown in Figure S2. $Mg^{2+}$-hexahydrates were placed near the DNA to neutralize its electrical charge of the DNA backbone.

We simulated two analogs of the dye conjugated DNA, one in an aqueous environment and another anchored in the lipid bilayer membrane leading to the data shown in Figure S3 and the Movies S1 and S2. To create the system in aqueous environment, the dye conjugated DNA molecule was solvated with TIP3P water molecules[20] using the Solvate plugin of VMD.[16] Potassium and chloride ions were added to produce 150 mM concentration of KCl in solution using the Autoionize plugin of VMD. Thus, assembled system measured 8 x 8 x 15 $nm^3$ and contained approximately 80,000 atoms. To create the membrane-anchored DNA system, we placed the dye conjugated DNA molecule in a pre-equilibrated patch of 1,2-dioleoyl-sn-glycero-3-phosphocholine (DOPC) lipid bilayer membrane such that the ATTO647N and $C_{12}$ spacer connecting it to the DNA span in the upper leaflet of the membrane. The lipid patch was generated using the CHARMM-GUI membrane builder[21] and pre-equilibrated for approximately 200 ns. Finally, we solvated the system with TIP3P water molecules[20] and added ions to produce a 150 mM concentration of KCl. Thus, the membrane-anchored system measured 10 x 10 x 15 $nm^3$ and contained approximately 130,000 atoms.

The assembled systems were subjected to energy minimization using the conjugate gradient method to remove the steric clashes between the solute and solvent. Following that, we equilibrated each system for 20 ns while harmonically restraining the phosphorus (P) atoms of DNA using a spring constant of 1 kcal $mol^{-1}$ $Å^{-2}$. Subsequently, we equilibrated the systems for additional 40 ns while maintaining the hydrogen bonds between the complementary base-pairs of DNA using the extrabond utility of NAMD. Finally, we removed all the restraints (except two P atoms of each DNA strand connecting the DNA to the origami plate) and performed approximately 1 μs long production simulations of systems using a constant number of atoms (N), pressure (P = 1 bar) and temperature (T = 300 K), the NPT ensemble. Two sets of simulations were carried for each design to improve sampling of the conformational space.

The simulation results presented in Figure S3b for the system without a membrane unravel a very close and stable state of the two fluorophores in the pink trajectory after ~0.75 μs. Inspecting Movie S1, melting of the dsDNA can be observed which leads to the dyes touching each other. Direct contact between dyes commonly yields complex photophysics with different intensity levels[22,23] that is not observed in our experiments. We therefore assign the DNA melting and the direct dye-dye contact to a force field artefact in the simulation which has been previously observed for CHARMM36 force fields as used in our system.[24] Therefore, for Figure 1f only data is included before the artefact is observed eliminating the prominent peak around 10 Å visible in Figure S3c. Also, for both the system with and without a lipid membrane, the first 0.2 μs of the simulation are excluded as this is approximately the time the system needs to equilibrate.

#### Double-membrane systems

To mimic the voltage bias created by a difference in the ionic concentration from inside to outside of a lipid vesicle in our simulations, we created a double membrane system having two identical patches of DOPC lipid bilayer membrane kept at a distance of 9 nm away from each other (distance between the center of the membranes) along the bilayer normal. We solvated the double DOPC membrane system using TIP3P water molecules[20] and added ions to produce 150 mM concentration of KCl. Thus, assembled double membrane system measured 11 x 11 x 20 $nm^3$ and contains 218,400 atoms.

The assembled system was subjected to energy minimization using the conjugate gradient method to remove the steric clashes between the solute and solvent. Following that, we equilibrated the double membrane system for 230 ns using the NPT ensemble. Towards the end of the equilibration, the distance between the center of the mass (CoM) of the individual bilayer along the bilayer normal stabilizes close to 8.8 nm. Averaging the dimensions of the simulation box from the last 20 ns of the NPT equilibration, we next performed simulation of the double membrane system in NVT ensemble. We created two other double membrane system named as 1 $K^+$ and -1 $K^+$, by shuffling one potassium ion from the inside chamber (the bulk water region around the center of the simulation box) to the outside chamber (the bulk water region at both ends of the simulation box) and *vice-versa*. Thus, we generated three double-membrane systems: 0 K+ system having exactly same number of potassium ions inside and outside, 1 $K^+$ system having two more potassium ions inside as compared to outside and -1 $K^+$ system having two fewer potassium ions inside as compared to the outside which is equivalent to a transmembrane potential of $\Delta\Psi=\pm180$ mV for this system's geometry. Finally, all systems were simulated using NVT ensemble with the exact same box dimension for approximately 300 ns. The data is presented in Figure S7.

#### Free energy calculations of dsDNA binding to double-membrane systems

Starting from a pre-equilibrated conformation of 21 base pair long dsDNA kept over a 10 x 7.1 $nm^2$ patch of DPhPE lipid membrane from our earlier study,[25] we created a DNA double membrane system by replicating another copy of the simulation cell along the bilayer normal. The DNA fragment is effectively infinite along its helical axis as both of the strands are connected to themselves across the periodic boundary (along the *y*-axis). We removed dsDNA and neutralizing counterions (42 potassium ions) from the outer chamber, which left us with the desired system having dsDNA only in the inner chamber of the double membrane and 150 mM concentration of KCl in solution (Figure S7a). Thus, assembled DNA double membrane system was measured 10 x 7.1 x 25 $nm^3$ and contains 181,628 atoms.

Both, the inner and outer chamber of this double membrane-DNA system are charge neutralized and we refer to this system as 0 $K^+$.

Next, we created two other systems, 4 $K^+$ and -4 $K^+$ by shuffling 4 potassium ions from the outer chamber to the inner chamber and *vice-versa* inducing a potential of $\Delta\Psi=\pm1.3$ V, a high enough value to observe statistically significant differences in the interactions between the DNA and the membrane without causing membrane electroporation. Thus, we have three systems with the same number of atoms but different potassium ions in inner and outer chambers. In 0 $K^+$, both the chambers are electrically charge-neutral, 4 $K^+$ system has 8$e^+$ charge in inner chamber as compared to the outer chamber and -4 $K^+$ system has 8 $e^+$ charge in the outer chamber as compared to the inner one due the shuffling of the potassium ions.

Twenty-one copies of each system were created by moving the CoM of dsDNA from 13 to 33 Å along the *z*-axis, the region shown using an arrow in Figure S7a. Note that Z =0 corresponds to the center of the simulation cell. Since the CoM of the upper membrane lies at $z$ = 63 Å, the distance of the CoM of dsDNA and upper membrane varies from 50 to 13 Å in respective copies of the simulation system. Replica exchange umbrella sampling simulations[26] were performed using the 1 Å sampling window for the distance between the CoM of dsDNA's and the upper membrane along the *z*-axis. A harmonic potential with the spring constant of 2.5 kcal/mol/ $Å^2$ was used the maintain the distance between dsDNA and the membrane in each window along the *z*-axis using colvars module[27] of NAMD. Each replica was run for approximately 120 ns. Weighted histogram analysis method (WHAM)[28] was used to subtract the effect of the harmonic potential and obtain the PMF profile. The first 5 ns of the simulation trajectories were excluded from the WHAM analysis.

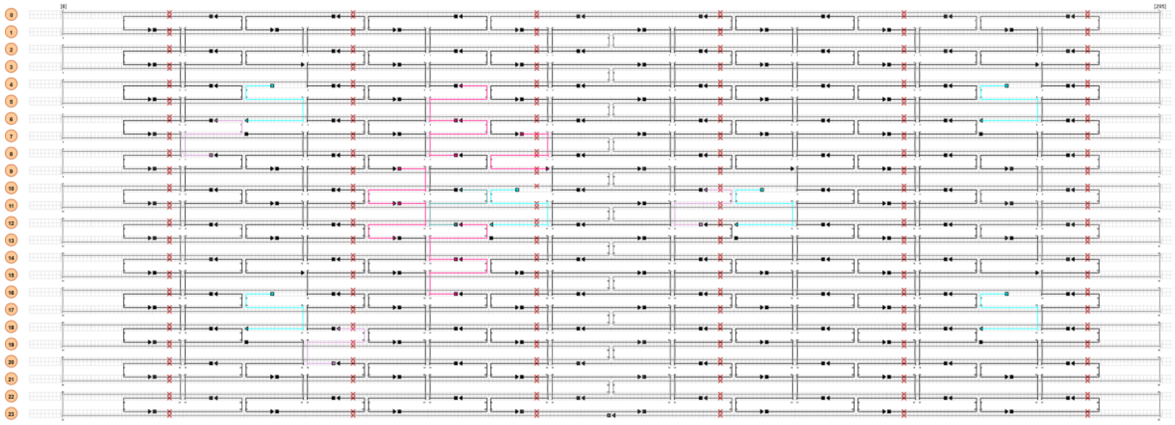

**Figure S1. Illustration of DNA origami design in CaDNAno[2].** The number of helices is shown in orange on the left and the nucleotide position described in the grid starting at position [8] and ending at position [295] (top). The scaffold strand is presented in dark grey, the unmodified staple strands in black, the biotinylated strands in blue carrying the modification on the 5' end, in light pink strands with cholesterol on the 3' end of a ssDNA spacer, in dark pink cholesterol attached via dsDNA on the 5' end and in cyan the voltage-sensing unit on the 5' end. The sequences of the strands and more details can be found in Table S1.

**Table S1. DNA oligonucleotides used as staple strands for the DNA origami voltage sensor.** The staple stands are listed with the 5' position x[y] with x as the helix number and y as the nucleotide number as described in Figure S1, the DNA sequence from the 5' to the 3' end, further modifications in the comment section and the company purchased from. Nucleotides in bold are not incorporated into the DNA origami, but protrude from the structure. For 12[111], either a or b are incorporated.

| 5' position | Sequence (5' to 3' end) | Comments | Company |
|---|---|---|---|
| / | **TATGAGTGTACACGATTGTTAAAA[SpC12][ATTO647N]** | Voltage sensor $C_{12}$+$C_6$, binds to 12[111]a strand [ATTO647N]: modification on amino-C6 linker [SpC12]: $C_{12}$ spacer | biomers.net GmbH |
| / | **[ATTO647N]AAATAACAATCGTGTACACTCATA** | Voltage sensor $C_{12}$, binds to 12[111]b strand [ATTO647N]: modification on amino-$C_{12}$ linker | biomers.net GmbH |
| 12[111] a | **TAACA[ATTO532]ATCGTGTACACTCATA**TAAATCATATAAC CTGTTTAGCTAACCTTTAA | Binding site for voltage sensor $C_{12}$+$C_6$ [ATTO532]: modification on amino-$C_6$-dT linker | Eurofins Genomics GmbH |
| 12[111] b | TAAATCATATAACCTGTTTAGCTAACCTTTAA**TATGAGTGTAC ACGAT[ATTO532]TGTTA** | Binding site for voltage sensor $C_{12}$ [ATTO532]: modification on amino-$C_6$-dT linker | Eurofins Genomics GmbH |
| 20[79] | TTCCAGTCGTAATCATGGTCATAAAAGGGG**AAAAAAAAAAA AAAAAAAAA [Chol]** | [Chol]: Cholesterol-TEG modification | Integrated DNA Technologies, Inc. |
| 8[47] | ATCCCCCTATACCACATTCAACTAGAAAAATC**AAAAAAAAAA AAAAAAAAAA [Chol]** | [Chol]: Cholesterol-TEG modification | Integrated DNA Technologies, Inc. |
| 12[175] | TTTTATTTAAGCAAATCAGATATTTTTTGT**AAAAAAAAAAAA AAAAAAAA [Chol]** | [Chol]: Cholesterol-TEG modification | Integrated DNA Technologies, Inc. |
| / | **GTGATGTAGGTGGTAGAGGA [Chol]** | Cholesterol strand, binds to multiple binding sites depicted below [Chol]: Cholesterol-TEG modification | Integrated DNA Technologies, Inc. |
| 11[96] | **TCCTCTACCACCTACATCAC**AATGGTCAACAGGCAAGGCAA AGAGTAATGTG | Cholesterol binding site | Eurofins Genomics GmbH |
| 14[111] | **TCCTCTACCACCTACATCAC**GAGGGTAGGATTCAAAAGGGT GAGACATCCAA | Cholesterol binding site | Eurofins Genomics GmbH |
| 6[111] | **TCCTCTACCACCTACATCAC**ATTACCTTTGAATAAGGCTTGC CCAAATCCGC | Cholesterol binding site | Eurofins Genomics GmbH |

| 5' position | Sequence (5' to 3' end) | Comments | Company |
|---|---|---|---|
| 7[128] | **TCCTCTACCACCTACATCAC**AGACGACAAAGAAGTTTTGCC ATAATTCGAGCTTCAA | Cholesterol binding site | Eurofins Genomics GmbH |
| 8[111] | **TCCTCTACCACCTACATCAC**AATAGTAAACACTATCATAACC CTCATTGTGA | Cholesterol binding site | Eurofins Genomics GmbH |
| 16[111] | **TCCTCTACCACCTACATCAC**TGTAGCCATTAAAATTCGCATT AAATGCCGGA | Cholesterol binding site | Eurofins Genomics GmbH |
| 9[96] | **TCCTCTACCACCTACATCAC**CGAAAGACTTTGATAAGAGGT CATATTTCGCA | Cholesterol binding site | Eurofins Genomics GmbH |
| 21[64] | GCCCTTCAGAGTCCACTATTAAAGGGTGCCGT | | Eurofins Genomics GmbH |
| 1[160] | TTAGGATTGGCTGAGACTCCTCAATAACCGAT | | Eurofins Genomics GmbH |
| 15[96] | ATATTTTGGCTTTCATCAACATTATCCAGCCA | | Eurofins Genomics GmbH |
| 22[271] | CAGAAGATTAGATAATACATTTGTCGACAA | | Eurofins Genomics GmbH |
| 5[128] | AACACCAAATTTCAACTTTAATCGTTTACC | | Eurofins Genomics GmbH |
| 20[47] | TTAATGAACTAGAGGATCCCCGGGGGGTAACG | | Eurofins Genomics GmbH |
| 18[79] | GATGTGCTTCAGGAAGATCGCACAATGTGA | | Eurofins Genomics GmbH |
| 12[207] | GTACCGCAATTCTAAGAACGCGAGTATTATTT | | Eurofins Genomics GmbH |
| 4[271] | AAATCACCTTCCAGTAAGCGTCAGTAATAA | | Eurofins Genomics GmbH |
| 22[207] | AGCCAGCAATTGAGGAAGGTTATCATCATTTT | | Eurofins Genomics GmbH |
| 8[207] | AAGGAAACATAAAGGTGGCAACATTATCACCG | | Eurofins Genomics GmbH |
| 15[128] | TAAATCAAAATAATTCGCGTCTCGGAAACC | | Eurofins Genomics GmbH |
| 6[271] | ACCGATTGTCGGCATTTTCGGTCATAATCA | | Eurofins Genomics GmbH |
| 0[79] | ACAACTTTCAACAGTTTCAGCGGATGTATCGG | | Eurofins Genomics GmbH |
| 11[224] | GCGAACCTCCAAGAACGGGTATGACAATAA | | Eurofins Genomics GmbH |
| 8[239] | AAGTAAGCAGACACCACGGAATAATATTGACG | | Eurofins Genomics GmbH |
| 19[96] | CTGTGTGATTGCGTTGCGCTCACTAGAGTTGC | | Eurofins Genomics GmbH |
| 15[224] | CCTAAATCAAAATCATAGGTCTAAACAGTA | | Eurofins Genomics GmbH |
| 1[192] | GCGGATAACCTATTATTCTGAAACAGACGATT | | Eurofins Genomics GmbH |
| 5[160] | GCAAGGCCTCACCAGTAGCACCATGGGCTTGA | | Eurofins Genomics GmbH |
| 12[239] | CTTATCATTCCCGACTTGCGGGAGCCTAATTT | | Eurofins Genomics GmbH |
| 2[111] | AAGGCCGCTGATACCGATAGTTGCGACGTTAG | | Eurofins Genomics GmbH |
| 19[248] | CGTAAAACAGAAATAAAAATCCTTTGCCCGAAAGATTAGA | | Eurofins Genomics GmbH |
| 0[111] | TAAATGAATTTTCTGTATGGGATTAATTTCTT | | Eurofins Genomics GmbH |
| 5[32] | CATCAAGTAAAACGAACTAACGAGTTGAGA | | Eurofins Genomics GmbH |
| 20[207] | GCGGAACATCTGAATAATGGAAGGTACAAAAT | | Eurofins Genomics GmbH |
| 12[143] | TTCTACTACGCGAGCTGAAAAGGTTACCGCGC | | Eurofins Genomics GmbH |
| 8[143] | CTTTTGCAGATAAAAACCAAAATAAAGACTCC | | Eurofins Genomics GmbH |
| 2[143] | ATATTCGGAACCATCGCCCACGCAGAGAAGGA | | Eurofins Genomics GmbH |
| 21[224] | CTTTAGGGCCTGCAACAGTGCCAATACGTG | | Eurofins Genomics GmbH |
| 14[239] | AGTATAAAGTTCAGCTAATGCAGATGTCTTTC | | Eurofins Genomics GmbH |
| 20[111] | CACATTAAAATTGTTATCCGCTCATGCGGGCC | | Eurofins Genomics GmbH |
| 3[128] | AGCGCGATGATAAATTGTGTCGTGACGAGA | | Eurofins Genomics GmbH |
| 17[128] | AGGCAAAGGGAAGGGCGATCGGCAATTCCA | | Eurofins Genomics GmbH |
| 17[160] | AGAAAACAAAGAAGATGATGAAACAGGCTGCG | | Eurofins Genomics GmbH |
| 2[207] | TTTCGGAAGTGCCGTCGAGAGGGTGAGTTTCG | | Eurofins Genomics GmbH |
| 22[79] | TGGAACAACCGCCTGGCCCTGAGGCCCGCT | | Eurofins Genomics GmbH |
| 10[47] | CTGTAGCTTGACTATTATAGTCAGTTCATTGA | | Eurofins Genomics GmbH |
| 16[207] | ACCTTTTTATTTTAGTTAATTTCATAGGGCTT | | Eurofins Genomics GmbH |

| 5' position | Sequence (5' to 3' end) | Comments | Company |
|---|---|---|---|
| 9[224] | AAAGTCACAAAATAAACAGCCAGCGTTTTA | | Eurofins Genomics GmbH |
| 19[224] | CTACCATAGTTTGAGTAACATTTAAAATAT | | Eurofins Genomics GmbH |
| 3[96] | ACACTCATCCATGTTACTTAGCCGAAAGCTGC | | Eurofins Genomics GmbH |
| 18[239] | CCTGATTGCAATATATGTGAGTGATCAATAGT | | Eurofins Genomics GmbH |
| 10[239] | GCCAGTTAGAGGGTAATTGAGCGCTTTAAGAA | | Eurofins Genomics GmbH |
| 3[32] | AATACGTTTGAAAGAGGACAGACTGACCTT | | Eurofins Genomics GmbH |
| 1[128] | TGACAACTCGCTGAGGCTTGCATTATACCA | | Eurofins Genomics GmbH |
| 16[47] | ACAAACGGAAAAGCCCCAAAAACACTGGAGCA | | Eurofins Genomics GmbH |
| 14[175] | CATGTAATAGAATATAAAGTACCAAGCCGT | | Eurofins Genomics GmbH |
| 17[192] | CATTTGAAGGCGAATTATTCATTTTTGTTTGG | | Eurofins Genomics GmbH |
| 19[56] | TACCGAGCTCGAATTCGGGAAACCTGTCGTGCAGCTGATT | | Eurofins Genomics GmbH |
| 6[111] | ATTACCTTTGAATAAGGCTTGCCCAAATCCGC | | Eurofins Genomics GmbH |
| 23[64] | AAAGCACTAAATCGGAACCCTAATCCAGTT | | Eurofins Genomics GmbH |
| 16[175] | TATAACTAACAAAGAACGCGAGAACGCCAA | | Eurofins Genomics GmbH |
| 7[56] | ATGCAGATACATAACGGGAATCGTCATAAATAAAGCAAAG | | Eurofins Genomics GmbH |
| 3[224] | TTAAAGCCAGAGCCGCCACCCTCGACAGAA | | Eurofins Genomics GmbH |
| 12[47] | TAAATCGGGATTCCCAATTCTGCGATATAATG | | Eurofins Genomics GmbH |
| 3[160] | TTGACAGGCCACCACCAGAGCCGCGATTTGTA | | Eurofins Genomics GmbH |
| 14[143] | CAACCGTTTCAAATCACCATCAATTCGAGCCA | | Eurofins Genomics GmbH |
| 6[79] | TTATACCACCAAATCAACGTAACGAACGAG | | Eurofins Genomics GmbH |
| 4[239] | GCCTCCCTCAGAATGGAAAGCGCAGTAACAGT | | Eurofins Genomics GmbH |
| 7[248] | GTTTATTTTGTCACAATCTTACCGAAGCCCTTTAATATCA | | Eurofins Genomics GmbH |
| 1[32] | AGGCTCCAGAGGCTTTGAGGACACGGGTAA | | Eurofins Genomics GmbH |
| 0[207] | TCACCAGTACAAACTACAACGCCTAGTACCAG | | Eurofins Genomics GmbH |
| 4[143] | TCATCGCCAACAAAGTACAACGGACGCCAGCA | | Eurofins Genomics GmbH |
| 16[271] | CTTAGATTTAAGGCGTTAAATAAAGCCTGT | | Eurofins Genomics GmbH |
| 13[96] | TAGGTAAACTATTTTTGAGAGATCAAACGTTA | | Eurofins Genomics GmbH |
| 21[192] | TGAAAGGAGCAAATGAAAAATCTAGAGATAGA | | Eurofins Genomics GmbH |
| 5[224] | TCAAGTTTCATTAAAGGTGAATATAAAAGA | | Eurofins Genomics GmbH |
| 14[111] | GAGGGTAGGATTCAAAAGGGTGAGACATCCAA | | Eurofins Genomics GmbH |
| 8[79] | AATACTGCCCAAAAGGAATTACGTGGCTCA | | Eurofins Genomics GmbH |
| 11[160] | CCAATAGCTCATCGTAGGAATCATGGCATCAA | | Eurofins Genomics GmbH |
| 4[111] | GACCTGCTCTTTGACCCCCAGCGAGGGAGTTA | | Eurofins Genomics GmbH |
| 2[175] | TATTAAGAAGCGGGGTTTTGCTCGTAGCAT | | Eurofins Genomics GmbH |
| 13[184] | GACAAAAGGTAAAGTAATCGCCATATTTAACAAAACTTTT | | Eurofins Genomics GmbH |
| 1[96] | AAACAGCTTTTTGCGGGATCGTCAACACTAAA | | Eurofins Genomics GmbH |
| 23[192] | ACCCTTCTGACCTGAAAGCGTAAGACGCTGAG | | Eurofins Genomics GmbH |
| 15[32] | TAATCAGCGGATTGACCGTAATCGTAACCG | | Eurofins Genomics GmbH |
| 18[175] | CTGAGCAAAAATTAATTACATTTTGGGTTA | | Eurofins Genomics GmbH |
| 6[175] | CAGCAAAAGGAAACGTCACCAATGAGCCGC | | Eurofins Genomics GmbH |
| 18[47] | CCAGGGTTGCCAGTTTGAGGGGACCCGTGGGA | | Eurofins Genomics GmbH |
| 6[143] | GATGGTTTGAACGAGTAGTAAATTTACCATTA | | Eurofins Genomics GmbH |
| 8[175] | ATACCCAACAGTATGTTAGCAAATTAGAGC | | Eurofins Genomics GmbH |

| 5' position | Sequence (5' to 3' end) | Comments | Company |
|---|---|---|---|
| 17[96] | GCTTTCCGATTACGCCAGCTGGCGGCTGTTTC | | Eurofins Genomics GmbH |
| 12[79] | AAATTAAGTTGACCATTAGATACTTTTGCG | | Eurofins Genomics GmbH |
| 8[271] | AATAGCTATCAATAGAAAATTCAACATTCA | | Eurofins Genomics GmbH |
| 15[192] | TCAAATATAACCTCCGGCTTAGGTAACAATTT | | Eurofins Genomics GmbH |
| 21[96] | AGCAAGCGTAGGGTTGAGTGTTGTAGGGAGCC | | Eurofins Genomics GmbH |
| 20[239] | ATTTTAAAATCAAAATTATTTGCACGGATTCG | | Eurofins Genomics GmbH |
| 7[224] | AACGCAAAGATAGCCGAACAAACCCTGAAC | | Eurofins Genomics GmbH |
| 20[79] | TTCCAGTCGTAATCATGGTCATAAAAGGGG | | Eurofins Genomics GmbH |
| 22[239] | TTAACACCAGCACTAACAACTAATCGTTATTA | | Eurofins Genomics GmbH |
| 18[111] | TCTTCGCTGCACCGCTTCTGGTGCGGCCTTCC | | Eurofins Genomics GmbH |
| 17[32] | TGCATCTTTCCCAGTCACGACGGCCTGCAG | | Eurofins Genomics GmbH |
| 4[175] | CACCAGAAAGGTTGAGGCAGGTCATGAAAG | | Eurofins Genomics GmbH |
| 13[120] | AAAGGCCGGAGACAGCTAGCTGATAAATTAATTTTTGT | | Eurofins Genomics GmbH |
| 10[175] | TTAACGTCTAACATAAAAACAGGTAACGGA | | Eurofins Genomics GmbH |
| 21[256] | GCCGTCAAAAAACAGAGGTGAGGCCTATTAGT | | Eurofins Genomics GmbH |
| 4[47] | GACCAACTAATGCCACTACGAAGGGGGTAGCA | | Eurofins Genomics GmbH |
| 6[239] | GAAATTATTGCCTTTAGCGTCAGACCGGAACC | | Eurofins Genomics GmbH |
| 18[271] | CTTTTACAAAATCGTCGCTATTAGCGATAG | | Eurofins Genomics GmbH |
| 14[79] | GCTATCAGAAATGCAATGCCTGAATTAGCA | | Eurofins Genomics GmbH |
| 2[47] | ACGGCTACAAAAGGAGCCTTTAATGTGAGAAT | | Eurofins Genomics GmbH |
| 21[128] | GCGAAAAATCCCTTATAAATCAAGCCGGCG | | Eurofins Genomics GmbH |
| 22[175] | ACCTTGCTTGGTCAGTTGGCAAAGAGCGGA | | Eurofins Genomics GmbH |
| 8[47] | ATCCCCCTATACCACATTCAACTAGAAAAATC | | Eurofins Genomics GmbH |
| 1[64] | TTTATCAGGACAGCATCGGAACGACACCAACCTAAAACGA | | Eurofins Genomics GmbH |
| 17[224] | CATAAATCTTTGAATACCAAGTGTTAGAAC | | Eurofins Genomics GmbH |
| 16[143] | GCCATCAAGCTCATTTTTTAACCACAAATCCA | | Eurofins Genomics GmbH |
| 4[207] | CCACCCTCTATTCACAAACAAATACCTGCCTA | | Eurofins Genomics GmbH |
| 2[79] | CAGCGAAACTTGCTTTCGAGGTGTTGCTAA | | Eurofins Genomics GmbH |
| 19[128] | CACAACAGGTGCCTAATGAGTGCCCAGCAG | | Eurofins Genomics GmbH |
| 19[192] | ATTATACTAAGAAACCACCAGAAGTCAACAGT | | Eurofins Genomics GmbH |
| 16[239] | GAATTTATTTAATGGTTTGAAATATTCTTACC | | Eurofins Genomics GmbH |
| 22[111] | GCCCGAGAGTCCACGCTGGTTTGCAGCTAACT | | Eurofins Genomics GmbH |
| 6[207] | TCACCGACGCACCGTAATCAGTAGCAGAACCG | | Eurofins Genomics GmbH |
| 13[224] | ACAACATGCCAACGCTCAACAGTCTTCTGA | | Eurofins Genomics GmbH |
| 20[175] | ATTATCATTCAATATAATCCTGACAATTAC | | Eurofins Genomics GmbH |
| 18[207] | CGCGCAGATTACCTTTTTTAATGGGAGAGACT | | Eurofins Genomics GmbH |
| 12[175] | TTTTATTTAAGCAAATCAGATATTTTTTGT | | Eurofins Genomics GmbH |
| 10[207] | ATCCCAATGAGAATTAACTGAACAGTTACCAG | | Eurofins Genomics GmbH |
| 14[207] | AATTGAGAATTCTGTCCAGACGACTAAACCAA | | Eurofins Genomics GmbH |
| 13[64] | TATATTTTGTCATTGCCTGAGAGTGGAAGATTGTATAAGC | | Eurofins Genomics GmbH |
| 7[128] | AGACGACAAAGAAGTTTTGCCATAATTCGAGCTTCAA | | Eurofins Genomics GmbH |
| 7[160] | TTATTACGAAGAACTGGCATGATTGCGAGAGG | | Eurofins Genomics GmbH |
| 23[96] | CCCGATTTAGAGCTTGACGGGGAAAAAGAATA | | Eurofins Genomics GmbH |

| 5’ position | Sequence (5’ to 3’ end) | Comments | Company |
|---|---|---|---|
| 3[192] | GGCCTTGAAGAGCCACCACCCTCAGAAACCAT | | Eurofins Genomics GmbH |
| 4[79] | GCGCAGACAAGAGGCAAAAGAATCCCTCAG | | Eurofins Genomics GmbH |
| 15[160] | ATCGCAAGTATGTAAATGCTGATGATAGGAAC | | Eurofins Genomics GmbH |
| 5[96] | TCATTCAGATGCGATTTTAAGAACAGGCATAG | | Eurofins Genomics GmbH |
| 18[143] | CAACTGTTGCGCCATTCGCCATTCAAACATCA | | Eurofins Genomics GmbH |
| 20[271] | CTCGTATTAGAAATTGCGTAGATACAGTAC | | Eurofins Genomics GmbH |
| 1[256] | CAGGAGGTGGGGTCAGTGCCTTGAGTCTCTGAATTTACCG | | Eurofins Genomics GmbH |
| 13[160] | GTAATAAGTTAGGCAGAGGCATTTATGATATT | | Eurofins Genomics GmbH |
| 1[224] | GTATAGCAAACAGTTAATGCCCAATCCTCA | | Eurofins Genomics GmbH |
| 5[192] | CGATAGCATTGAGCCATTTGGGAACGTAGAAA | | Eurofins Genomics GmbH |
| 7[192] | ATACATACCGAGGAAACGCAATAAGAAGCGCATTAGACGG | | Eurofins Genomics GmbH |
| 10[111] | TTGCTCCTTTCAAATATCGCGTTTGAGGGGGT | | Eurofins Genomics GmbH |
| 13[256] | GTTTATCAATATGCGTTATACAAACCGACCGTGTGATAAA | | Eurofins Genomics GmbH |
| 6[47] | TACGTTAAAGTAATCTTGACAAGAACCGAACT | | Eurofins Genomics GmbH |
| 16[79] | GCGAGTAAAAATATTTAAATTGTTACAAAG | | Eurofins Genomics GmbH |
| 23[256] | CTTTAATGCGCGAACTGATAGCCCCACCAG | | Eurofins Genomics GmbH |
| 23[224] | GCACAGACAATATTTTTGAATGGGGTCAGTA | | Eurofins Genomics GmbH |
| 10[127] | TAGAGAGTTATTTTCATTTGGGGATAGTAGTAGCATTA | | Eurofins Genomics GmbH |
| 4[63] | ATAAGGGAACCGGATATTCATTACGTCAGGACGTTGGGAA | | Eurofins Genomics GmbH |
| 16[63] | CGGATTCTGACGACAGTATCGGCCGCAAGGCGATTAAGTT | | Eurofins Genomics GmbH |
| 10[191] | GAAACGATAGAAGGCTTATCCGGTCTCATCGAGAACAAGC | | Eurofins Genomics GmbH |
| 16[255] | GAGAAGAGATAACCTTGCTTCTGTTCGGGAGAAACAATAA | | Eurofins Genomics GmbH |
| 4[255] | AGCCACCACTGTAGCGCGTTTTCAAGGGAGGGAAGGTAAA | | Eurofins Genomics GmbH |
| 10[79] | GATGGCTTATCAAAAAGATTAAGAGCGTCC | | Eurofins Genomics GmbH |
| 0[47] | AGAAAGGAACAACTAAAGGAATTCAAAAAAA | | Eurofins Genomics GmbH |
| 10[271] | ACGCTAACACCCACAAGAATTGAAAATAGC | | Eurofins Genomics GmbH |
| 2[271] | GTTTTAACTTAGTACCGCCACCCAGAGCCA | | Eurofins Genomics GmbH |
| 14[271] | TTAGTATCACAATAGATAAGTCCACGAGCA | | Eurofins Genomics GmbH |
| 9[160] | AGAGAGAAAAAAATGAAAATAGCAAGCAAACT | | Eurofins Genomics GmbH |
| 12[271] | TGTAGAAATCAAGATTAGTTGCTCTTACCA | | Eurofins Genomics GmbH |
| 23[160] | TAAAAGGGACATTCTGGCCAACAAAGCATC | | Eurofins Genomics GmbH |
| 21[160] | TCAATATCGAACCTCAAATATCAATTCCGAAA | | Eurofins Genomics GmbH |
| 19[160] | GCAATTCACATATTCCTGATTATCAAAGTGTA | | Eurofins Genomics GmbH |
| 10[143] | CCAACAGGAGCGAACCAGACCGGAGCCTTTAC | | Eurofins Genomics GmbH |
| 23[32] | CAAATCAAGTTTTTTGGGGTCGAAACGTGGA | | Eurofins Genomics GmbH |
| 22[143] | TCGGCAAATCCTGTTTGATGGTGGACCCTCAA | | Eurofins Genomics GmbH |
| 0[175] | TCCACAGACAGCCCTCATAGTTAGCGTAACGA | | Eurofins Genomics GmbH |
| 0[143] | TCTAAAGTTTTGTCGTCTTTCCAGCCGACAA | | Eurofins Genomics GmbH |
| 20[143] | AAGCCTGGTACGAGCCGGAAGCATAGATGATG | | Eurofins Genomics GmbH |
| 2[239] | GCCCGTATCCGGAATAGGTGTATCAGCCCAAT | | Eurofins Genomics GmbH |
| 7[32] | TTTAGGACAAATGCTTTAAACAATCAGGTC | | Eurofins Genomics GmbH |
| 23[128] | AACGTGGCGAGAAAGGAAGGGAAACCAGTAA | | Eurofins Genomics GmbH |
| 21[32] | TTTTCACTCAAAGGGCGAAAAACCATCACC | | Eurofins Genomics GmbH |

| 5' position | Sequence (5' to 3' end) | Comments | Company |
|---|---|---|---|
| 14[47] | AACAAGAGGGATAAAAATTTTTAGCATAAAGC | | Eurofins Genomics GmbH |
| 13[32] | AACGCAAAATCGATGAACGGTACCGGTTGA | | Eurofins Genomics GmbH |
| 0[271] | CCACCCTCATTTTCAGGGATAGCAACCGTACT | | Eurofins Genomics GmbH |
| 9[256] | GAGAGATAGAGCGTCTTTCCAGAGGTTTTGAA | | Eurofins Genomics GmbH |
| 11[256] | GCCTTAAACCAATCAATAATCGGCACGCGCCT | | Eurofins Genomics GmbH |
| 0[239] | AGGAACCCATGTACCGTAACACTTGATATAA | | Eurofins Genomics GmbH |
| 9[32] | TTTACCCCAACATGTTTTAAATTTCCATAT | | Eurofins Genomics GmbH |
| 11[32] | AACAGTTTTGTACCAAAAACATTTTATTTC | | Eurofins Genomics GmbH |
| 22[47] | CTCCAACGCAGTGAGACGGGCAACCAGCTGCA | | Eurofins Genomics GmbH |
| 19[32] | GTCGACTTCGGCCAACGCGCGGGGTTTTTC | | Eurofins Genomics GmbH |
| 12[111] | TAAATCATATAACCTGTTTAGCTAACCTTTAA | | Eurofins Genomics GmbH |
| 7[96] | TAAGAGCAAATGTTTAGACTGGATAGGAAGCC | | Eurofins Genomics GmbH |
| 8[111] | AATAGTAAACACTATCATAACCCTCATTGTGA | | Eurofins Genomics GmbH |
| 16[111] | TGTAGCCATTAAAATTCGCATTAAATGCCGGA | | Eurofins Genomics GmbH |
| 9[96] | CGAAAGACTTTGATAAGAGGTCATATTTCGCA | | Eurofins Genomics GmbH |
| 11[64] | GATTTAGTCAATAAAGCCTCAGAGAACCCTCA | | Eurofins Genomics GmbH |
| 9[64] | CGGATTGCAGAGCTTAATTGCTGAAACGAGTA | | Eurofins Genomics GmbH |
| 11[96] | AATGGTCAACAGGCAAGGCAAAGAGTAATGTG | | Eurofins Genomics GmbH |

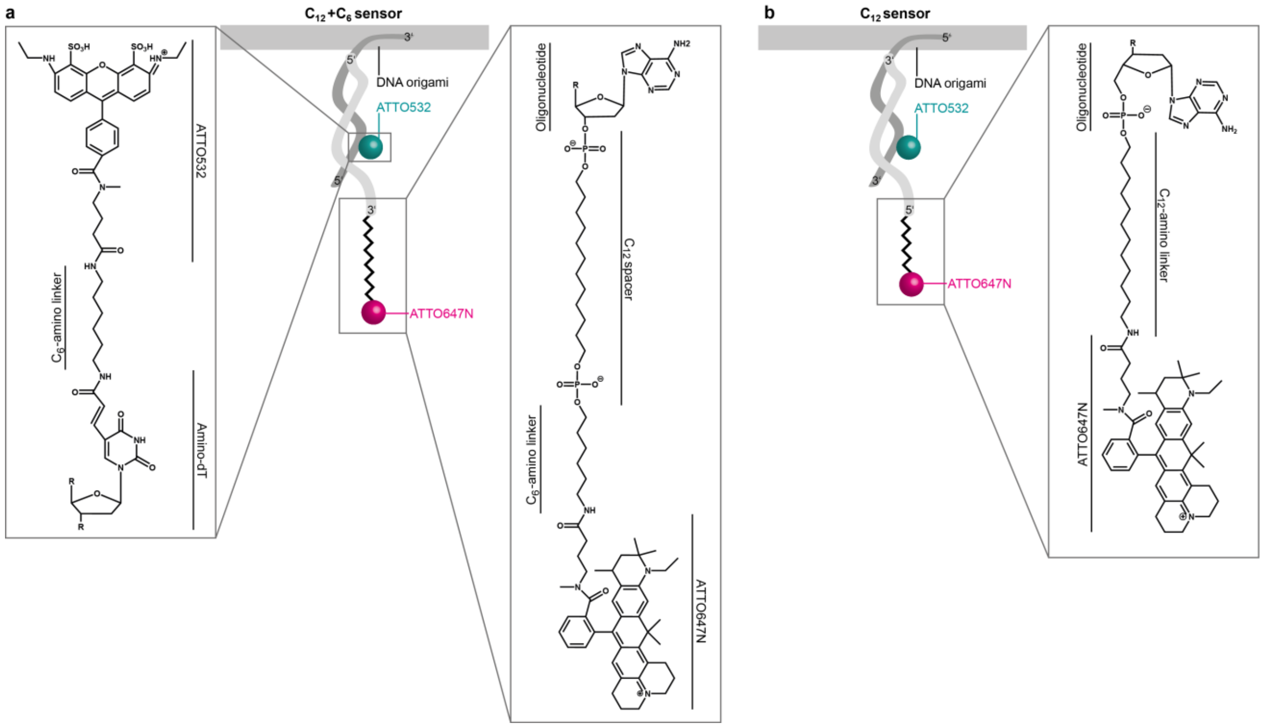


**Figure S2. Detailed sketch of the different voltage-sensor designs.** (a) For the $C_{12}$+$C_6$ sensor, a strand from the DNA origami is extended from the 5' end (dark grey) and carries an ATTO532 (cyan, zoom-in left for details). In light grey, the counter strand is shown with the 3' end carrying a carbon chain and an ATTO647N (pink). The right zoom-in shows how the ATTO647N is connected to the DNA's 3' end via a $C_{12}$ spacer and a $C_6$-amino linker. (b) For the $C_{12}$ sensor, the strand in the origami is extended on the 3' end (dark grey) and also carries the ATTO532 (cyan) which is connected as shown in the zoom-in in (a). The counter strand carries on the 5' end the carbon chain with the ATTO647N. In the zoom-in, the connection between the 5' end of the DNA and the ATTO647N via a $C_{12}$-amino linker is shown.

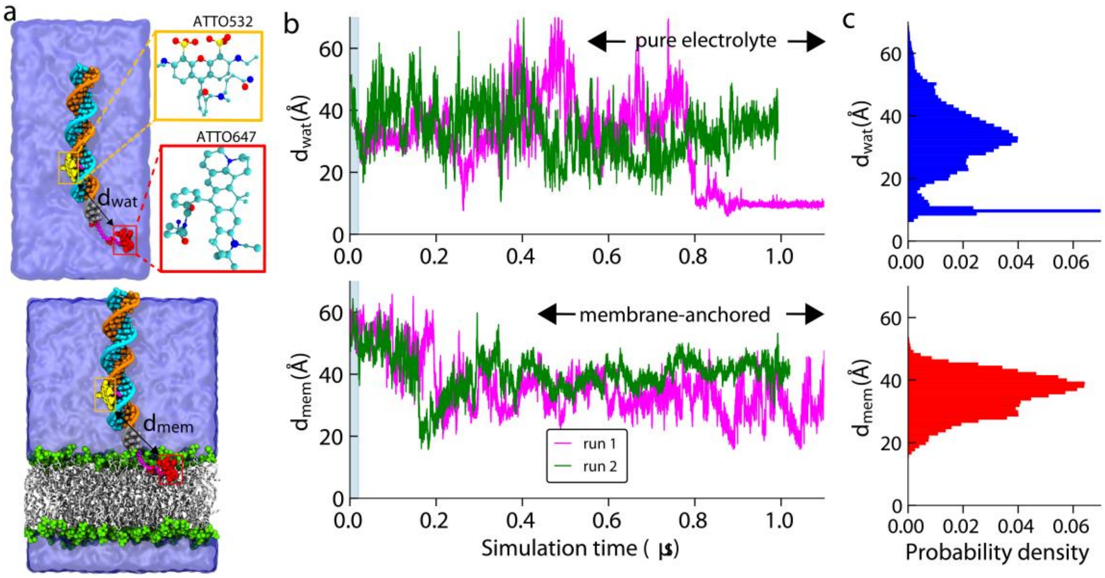


**Figure S3. Equilibrium MD simulation of dye conjugated dsDNA in aqueous and membrane anchored environment.** (a) A cut-away view of the all-atom models of 22-base pair long dsDNA conjugated to ATTO532 (yellow) and ATTO647N (red) dye molecules, solvated in the aqueous solution (top panel); anchored to lipid bilayer membrane and then solvated (bottom panel). The alkyl carbon chains ($C_6$ and $C_{12}$) connecting the dye molecules to the DNA (as shown in the chemical sketch in Figure S2) are shown in magenta. Complementary strands of the DNA are shown in turquoise and orange, the DOPC lipid head groups are shown as green spheres whereas the lipid tails are shown as white lines, water and ions are not shown for the sake of clarity. The enlarged atomic structures of the respective dye molecules are shown in yellow and red boxes. (b) Distance between the center of mass of the ATTO532 and ATTO647N dye molecules as a function of simulation time from two independent simulation runs in a box of water (top panel) and while anchored in lipid bilayer membrane (bottom panel). (c) The histogram of the distance between the dye molecules in water (top) and while anchored in membrane (bottom) excluding the first 200 ns from each simulation run.

**Table S2. Concentration of KCl and NaCl in the buffer inside and outside of the LUVs.** To build up a certain transmembrane potential ΔΨ with respect to the Nernst equation, the buffers used inside and outside of the LUVs were varied. The KCl gradient is responsible for the potential creation whereas NaCl is added to prevent osmotic pressure. The NaCl/KCl concentrations add up to buffers containing 5 mM Tris, 1 mM EDTA and 0.5 mM Trolox at pH 7 as previously described as the LUV buffer.

| | $c_{inside}$ [mM] | | $c_{outside}$ [mM] | |
|---|---|---|---|---|
| **ΔΨ [mV]** | **KCl** | **NaCl** | **KCl** | **NaCl** |
| -125 | 150 | 0 | 1.1 | 148.9 |
| -100 | 150 | 0 | 3 | 147 |
| -75 | 150 | 0 | 8 | 142 |
| -50 | 150 | 0 | 21 | 129 |
| -25 | 150 | 0 | 56 | 94 |
| 0 | 150 | 0 | 150 | 0 |
| 0 | 1 | 149 | 1 | 149 |
| 25 | 1 | 149 | 2.7 | 147.3 |
| 50 | 1 | 149 | 7 | 143 |
| 75 | 1 | 149 | 19 | 131 |
| 100 | 1 | 149 | 50 | 100 |
| 125 | 1 | 149 | 134 | 16 |

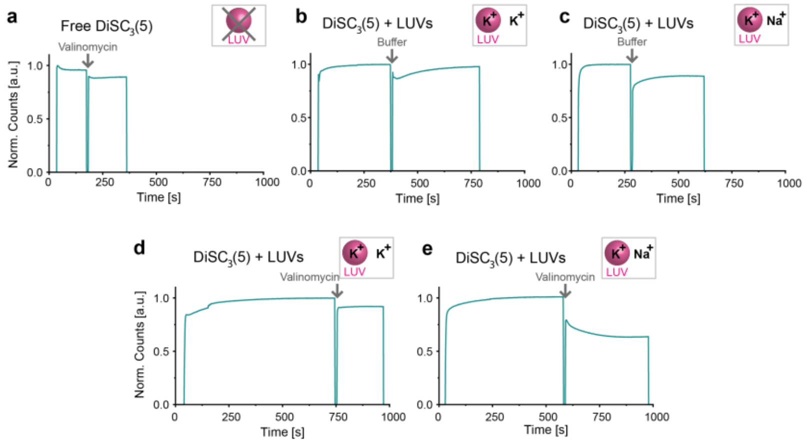


**Figure S4. Valinomycin bulk test.** On a spectrofluorometer, it was proven that a transmembrane potential ΔΨ is built up with the valinomycin assay using the voltage-sensitive dye DiSC$_3$(5). As a control, the DiSC$_3$(5) was investigated without LUVs in the cuvette and valinomycin was added to the solution (a). The intensity drop results from the dye sticking to the pipette tip while adding the solution. No further effect is observed. As another control, LUVs with potassium buffer inside and outside of the liposome (b) and inside potassium and outside sodium (c) were used and buffer was added to the samples. The intensity drop due to dye sticking to the pipette is observed as previously. Further, the intensity equilibrates after the buffer addition for both samples which is related to the homogeneous distribution after Brownian motion in the field of view. Next, valinomycin was added to LUVs with inside and outside potassium ions and buffer was injected (d). As for the other controls, an intensity drop due to dye sticking is observed, but no further effect. Only LUVs with inside potassium and outside sodium show a clear drop in intensity (e) which equilibrates after a while following Brownian motion. As more of the DiSC$_3$(5) is attracted to the charged membrane, contact quenching of the dyes take place and the intensity is reduced. For details see the chapter on "Valinomycin bulk assay" above.

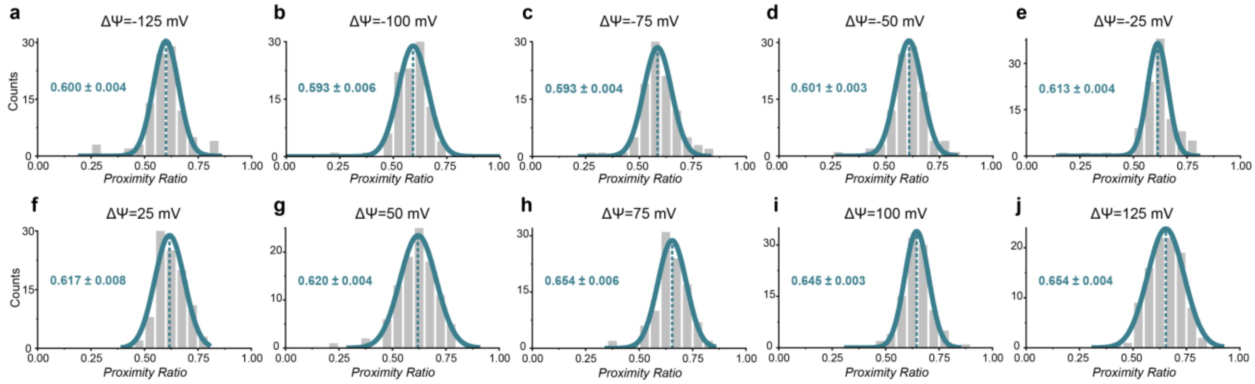


**Figure S5. *PR* distributions for all samples of the $C_{12}$+$C_6$ sensor with transmembrane potentials ΔΨ.** The samples were imaged after buffer exchange, addition of valinomycin and hence, the creation of a transmembrane potential of (a) ΔΨ=-125 mV, (b) ΔΨ=-100 mV, (c) ΔΨ=-75 mV, (d) ΔΨ=-50 mV, (e) ΔΨ=-25 mV, (f) ΔΨ=25 mV, (g) ΔΨ=50 mV, (h) ΔΨ=75 mV, (i) ΔΨ=100 mV and (j) ΔΨ=125 mV. The respective mean resulting from the Gauss fit as well as the standard error of the mean is given for each distribution. $N_{molecule}$: (a)-(j) 100.

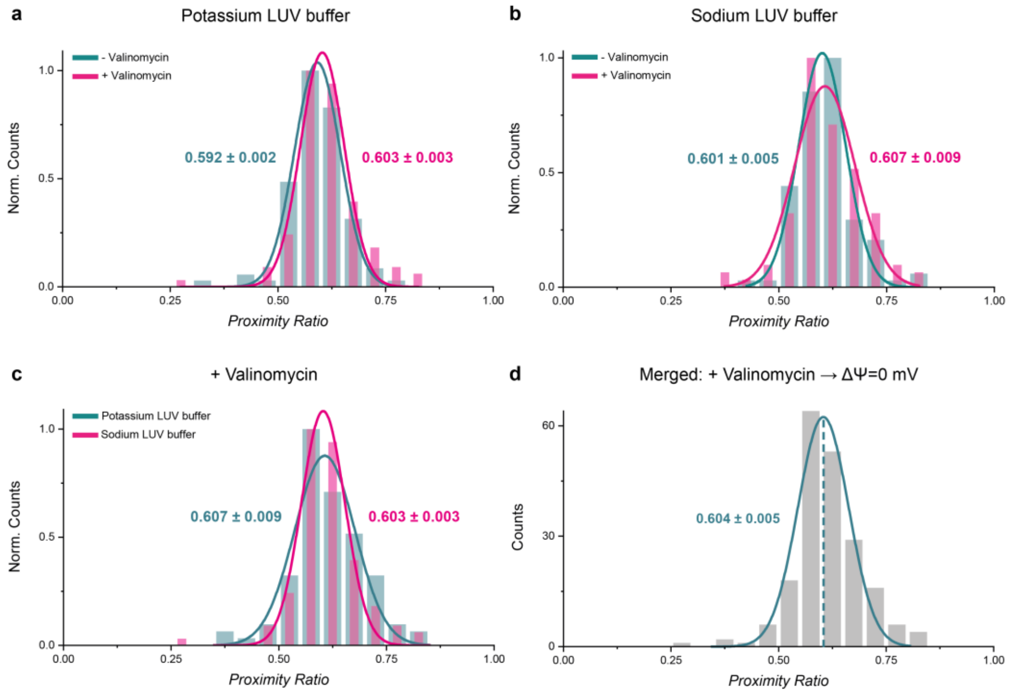


**Figure S6. *PR* distributions of various $C_{12}$+$C_6$ samples compared.** Control for ΔΨ=0 mV with potassium LUV buffer in cyan with the sample after the addition of valinomycin in pink (a), control for ΔΨ=0 mV with sodium LUV buffer in cyan with the sample after the addition of valinomycin in pink (b) and both ΔΨ=0 mV samples in potassium (cyan) and sodium (pink) LUV buffer with their respective mean from Gauss fitting and the standard error of the mean. All derived mean values are very similar which is why on the one hand a buffer effect on the *PR* value and on the other hand an effect of the valinomycin can be neglected. Therefore, the data for both samples with valinomycin (c) are merged and presented in (d) which in the following is the ΔΨ=0 mV sample. $N_{molecule}$: (a)-(c) 100 for each distribution, (d) 200.

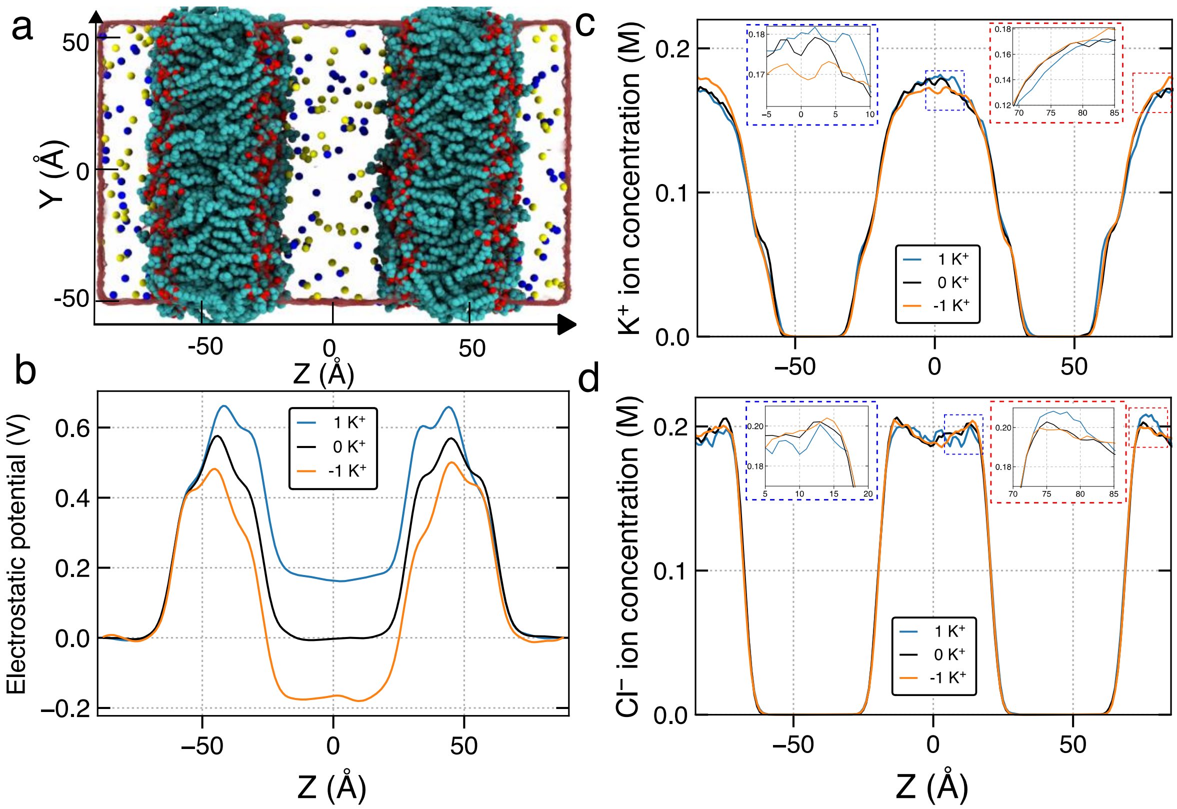


**Figure S7. Voltage bias created by shuffling of a single ion in double membrane systems.** (a) A representative snapshot of the all-atom model of double DOPC lipid bilayer membrane solvated in 150 mM solution of KCl. Non-hydrogen atoms of the lipid (DOPC) membrane are shown as blue (N), tan (P), red (O), and cyan (C) spheres. The transparent surface illustrates the volume occupied by the water molecules. The potassium and chloride ions are shown in blue and yellow spheres. (b) Average electrostatic potential profile of various systems along the *z*-axis. Local concentration of (c) potassium and (d) chloride ions along the lipid bilayer normal averaged over last 300 ns of NVT simulation. The images inside the close-in boxes shows the zoomed in region in the plots. The density was obtained by dividing the *z*-axis in the bins of 1 Å and then Savitzky Golay filter (window length 21 with a polynomial of degree 2) is used for smoothening of the data.

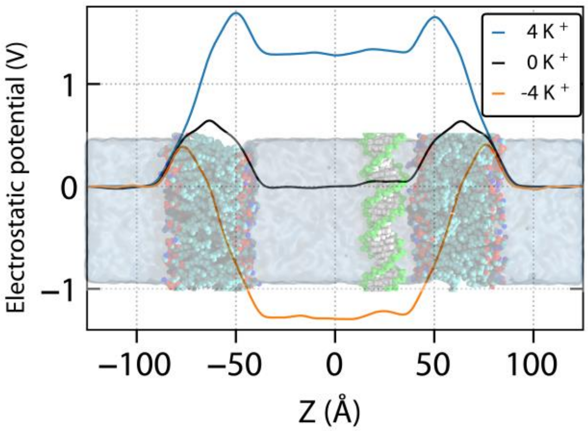


**Figure S8. Average electrostatic profile of the 0 K⁺, 4 K⁺ and -4 K⁺ systems along the bilayer normal during the REUS MD simulations.**

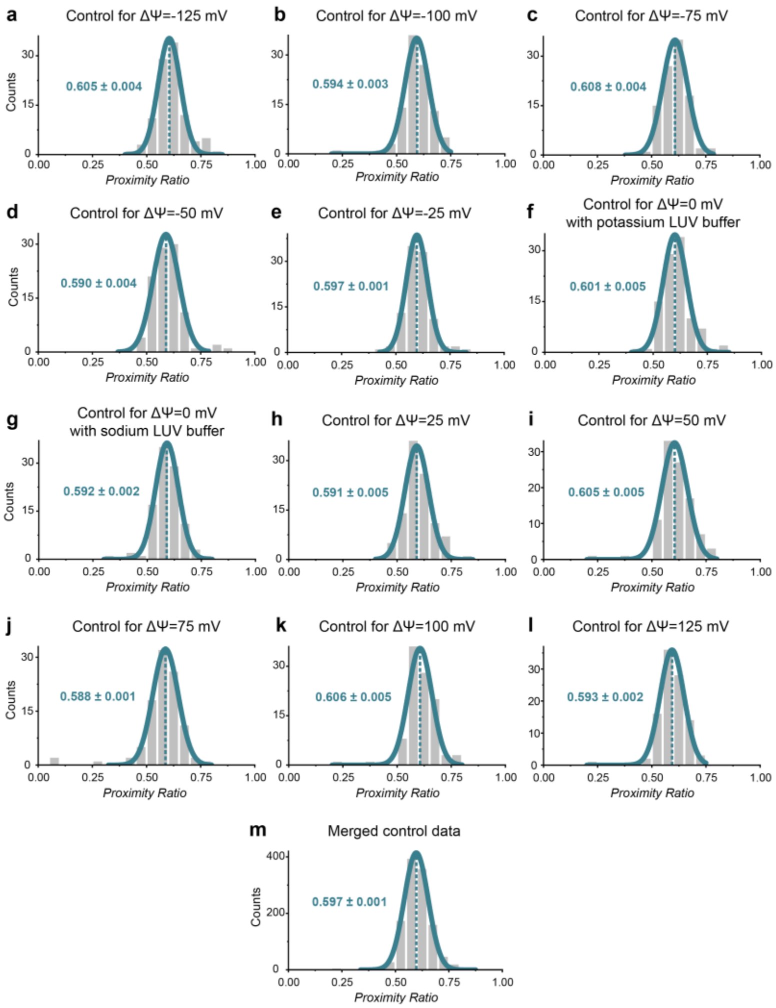


**Figure S9. *PR* distributions for all control samples of the $C_{12}$+$C_6$ sensor.** The samples were imaged before buffer exchange and transmembrane potential creation via valinomycin and the plots show the distributions with their respective mean and standard error of the mean from the Gauss fit. Control sample resulting in (a) ΔΨ=-125 mV, (b) ΔΨ=-100 mV, (c) ΔΨ=-75 mV, (d) ΔΨ=-50 mV, (e) ΔΨ=-25 mV, (f) ΔΨ=0 mV with potassium LUV buffer, (g) ΔΨ=0 mV with sodium LUV buffer, (h) ΔΨ=25 mV, (i) ΔΨ=50 mV, (j) ΔΨ=75 mV, (k) ΔΨ=100 mV, (l) ΔΨ=125 mV. As all of the samples show similar mean values, their data is merged and plotted together in (m) and further referred to as the sample before the valinomycin addition. $N_{molecule}$: (a)-(l) 100, (m) 1200.

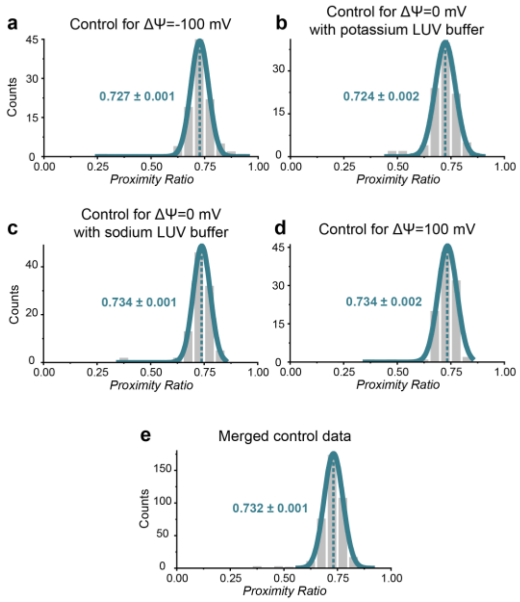


**Figure S10. *PR* distributions for all control samples of the $C_{12}$ sensor.** The samples were imaged before buffer exchange and transmembrane potential creation via valinomycin and the plots show the distributions with their respective mean and standard error of the mean from the Gauss fit. Control sample resulting in (a) ΔΨ=-100 mV, (b) ΔΨ=0 mV with potassium LUV buffer, (c) ΔΨ=0 mV with sodium LUV buffer and (d) ΔΨ=125 mV. As all of the samples show similar mean values, their data is merged and plotted together in (e) and further referred to as the sample before the valinomycin addition. $N_{\mathrm{molecule}}$: (a)-(d) 100, (e) 400.

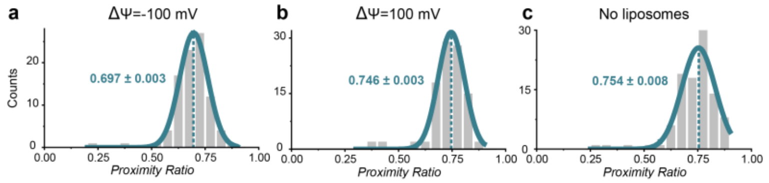


**Figure S11. *PR* distributions for samples of the $C_{12}$ sensor with transmembrane potentials ΔΨ and without liposomes.** For the creation of transmembrane potentials of (a) ΔΨ=-100 mV and (b) ΔΨ=100 mV, the outside buffer was exchanged and valinomycin was added to create the potentials. (c) *PR* distribution for sample without liposomes. The respective mean resulting from the Gauss fit as well as the standard error of the mean is given for each distribution. $N_{molecule}$: (a) 91, (b)-(c) 100.

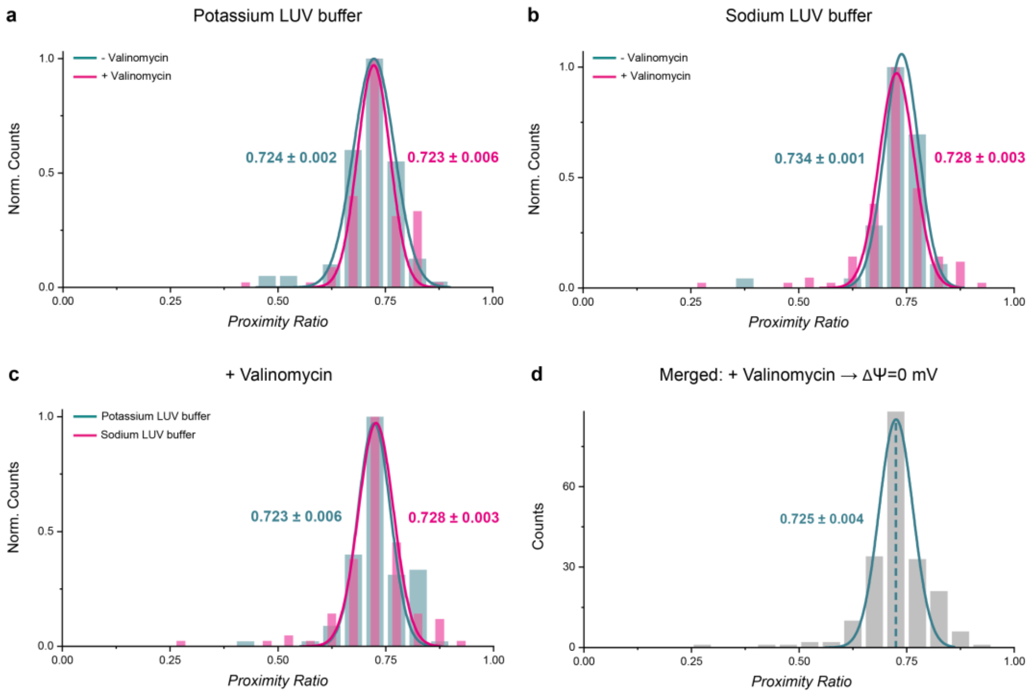


**Figure S12. *PR* distributions of various $C_{12}$ samples compared.** Control for ΔΨ=0 mV with potassium LUV buffer in cyan with the sample after the addition of valinomycin in pink (a), control for ΔΨ=0 mV with sodium LUV buffer in cyan with the sample after the addition of valinomycin in pink (b) and both ΔΨ=0 mV samples in potassium (cyan) and sodium (pink) LUV buffer with their respective mean from Gauss fitting and the standard error of the mean. All derived mean values are very similar which is why on the one hand a buffer effect on the *PR* and on the other hand an effect of the valinomycin can be neglected. Therefore, the data for both samples with valinomycin (c) are merged and presented in (d) which in the following is the ΔΨ=0 mV sample. $N_{molecule}$: (a) cyan: 100, pink: 99, (b) each 100, (c) cyan: 99, pink: 100, (d) 199.

*Captions to Supplementary Movies*

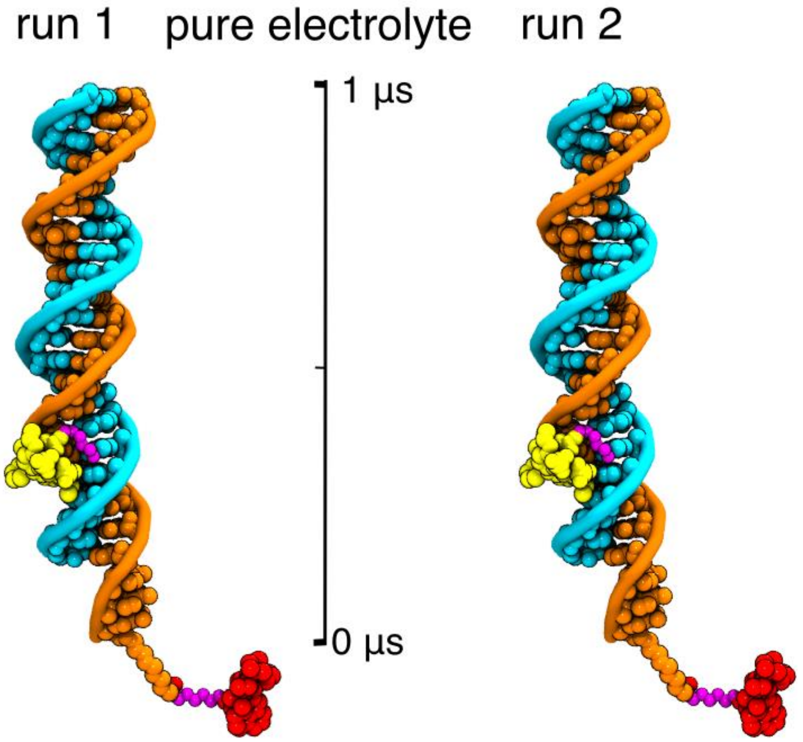


**Supplementary Movie 1.** All-atom molecular dynamics simulation of ATTO532 (yellow) and ATTO647N (red) dye molecules conjugated to dsDNA in purely aqueous solution. The movie illustrates MD trajectories of two independent simulation runs (run 1 and run 2) starting from the same initial configuration. Water and ions are not shown for clarity.

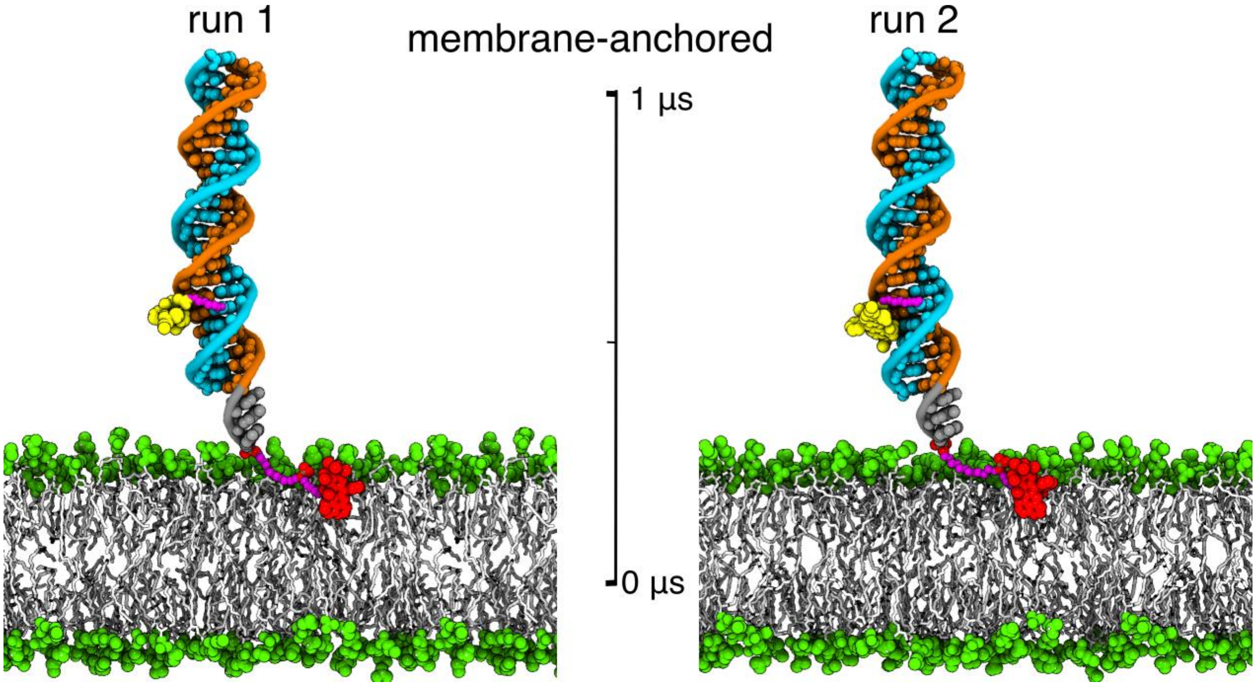


**Supplementary Movie 2.** All-atom molecular dynamics simulation of ATTO532 (yellow) and ATTO647N (red) dye molecules conjugated to dsDNA anchored in DOPC lipid bilayer membrane. The movie illustrates MD trajectories of two independent simulation runs (run 1 and run 2) starting from the same initial configuration. Water and ions are not shown for clarity.

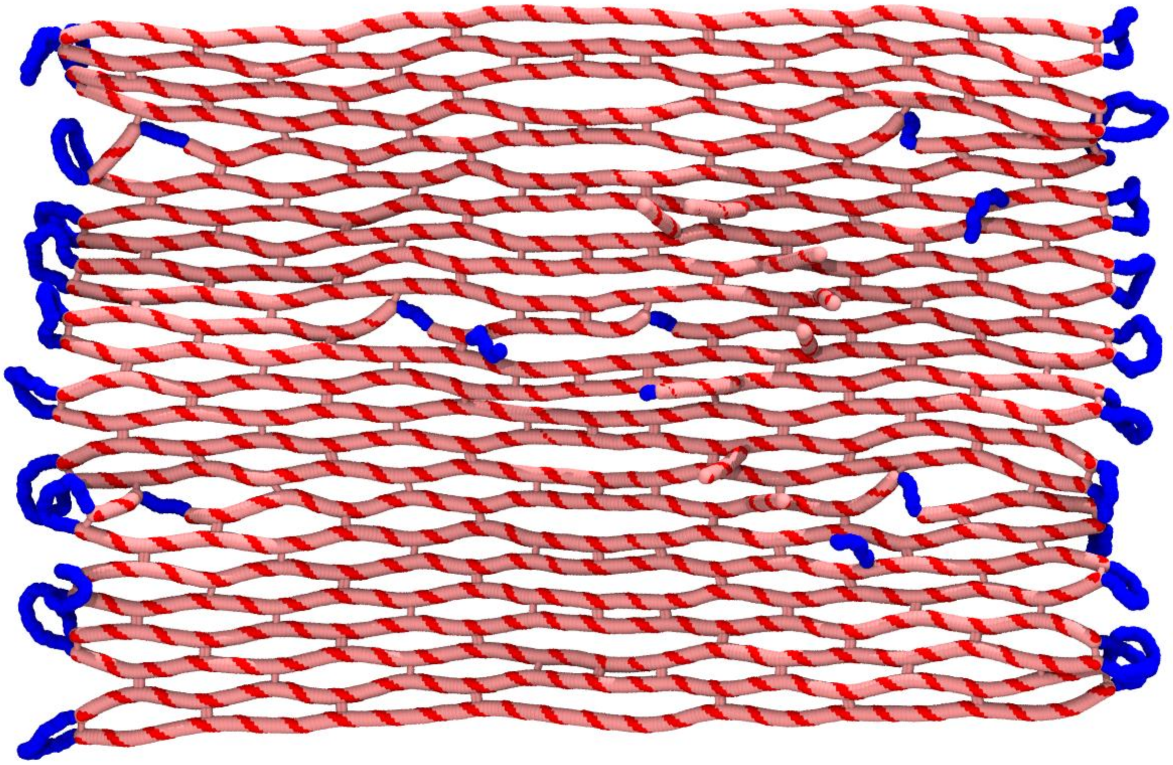

**Supplementary Movie 3. A typical mrDNA[18] simulation of the DNA origami plate designed as a sensor of transmembrane potentials**. The coarse-grained simulation starts with the caDNAno design of the origami plate which is first mapped into a 5-bp/bead model followed by a 1-bead/bp model. Finally, the all-atom model of the system was obtained by averaging equilibrated conformations in coarse-grained simulation. At the end, the all-atom model was simulated for couple of nanoseconds in vacuum using the network of elastic restraints.[29]
